# Hybrid origin of *Populus tomentosa* Carr. identified through genome sequencing and phylogenomic analysis

**DOI:** 10.1101/2020.04.07.030692

**Authors:** Xinmin An, Kai Gao, Zhong Chen, Juan Li, Xiong Yang, Xiaoyu Yang, Jing Zhou, Ting Guo, Tianyun Zhao, Sai Huang, Deyu Miao, Wasif Ullah Khan, Pian Rao, Meixia Ye, Bingqi Lei, Weihua Liao, Jia Wang, Lexiang Ji, Ying Li, Bing Guo, Nada Siddig Mustafa, Shanwen Li, Quanzheng Yun, Stephen R. Keller, Jianfeng Mao, Rengang Zhang, Steven H. Strauss

**Affiliations:** Beijing Advanced Innovation Center for Tree Breeding by Molecular Design, Beijing Forestry University, Beijing 100083, China; National Engineering Laboratory for Tree Breeding, College of Biological Sciences and Technology, Beijing Forestry University, Beijing 100083, China; Key Laboratory of Genetics and Breeding in Forest Trees and Ornamental Plants, MOE, College of Biological Sciences and Technology, Beijing Forestry University, Beijing 100083, China; Shanxi Academy of Forestry, Taiyuan 030012, China; Shandong Academy of Forestry, Jinan 250014, China; Ori-Gene Technology Co.,Ltd. Beijing 102206, China; Department of Plant Biology, University of Vermont, 111 Jeffords Hall, Burlington, VT 05405, USA; Department of Forest Ecosystems and Society, Oregon State University, Corvallis, OR 97331 USA

**Keywords:** *Populus tomentosa*, PacBio long-read sequencing, Genome assembly, Hybridization, Forest biotechnology

## Abstract

*Populus tomentosa* is widely distributed and cultivated in the Northern and Central China, where it is of great economic and ecological importance. However, the origin of *P. tomentosa* remains controversial. Here, we used a PacBio+Hi-C+Illumina strategy to sequence and assemble its 740.2 Mb (2n) genome. The assembly accounts for greater than 92.1% of the 800-megabase genome, comprises 38 chromosomes, and contains 59,124 annotated protein-coding genes. Phylogenomic analyses elucidated dynamic genome evolution events among its closely related white poplars, and revealed that *tomentosa* is comprised of two subgenomes, which we deomonstrate is likely to have resulted from hybridization between *Populus adenopoda* as the female, and *Populus alba* var. *pyramidalis* as the male, around 3.93 Mya. We also detected structural variations and allele-indels across genome. Our study presents a high quality and well assembled genome, unveils the origin of the widely distributed and planted *P. tomentosa*, and provides a powerful resource for comparative plant biology, breeding, and biotechnology.

## Introduction

The genomics revolution has spurred unprecedented growth in the sequencing and assembly of whole genomes in a wide variety of model and non-model organisms (Ellegren, 2014). While this has fueled the development of large genomic diversity panels for studies into the genetic basis of adaptive traits, reliance on a single well-assembled reference genome within a species or across a set of closely related congeners poses significant limitations on genetic and evolutionary inferences (Ballouz et al., 2019; Sherman and Salzberg, 2020). The challenge is particularly acute when working with large, structurally diverse, hybrid or heterozygous genomes, for which low coverage and biases in variant calling may result when mapping short read sequences against a divergent reference genome.

Long-lived perennial forest trees present unique challenges and opportunities for evolutionary genomics, due to abundant structural and nucleotide diversity within their genomes, even among close congeners. The genus *Populus* (poplars, cottonwoods, and aspens) has emerged as the leading model in tree ecological genomics and biotechnology, including development of the reference genome assembly for *Populus trichocarpa*–the first tree to undergo whole genome sequencing (Tuskan et al., 2006). In recent years, the whole genomes of *Populus euphratica, Populus pruinosa, Populus tremula* and *Populus tremuloides*, and hybrid 84K (*P. alba* x *P. tremula var. glandulosa*) have also been published (Lin et al., 2018; Ma et al., 2013; Qiu et al., 2019; Yang et al., 2017). However, high genetic heterozygosity and short read lengths limit the quality of these genome assemblies, which remain highly fragmented into thousands of scaffolds (Ambardar et al., 2016).

The availability of multiple highly contiguous, well-assembled *Populus* reference genomes would greatly facilitate accurate inferences of synteny, recombination, and chromosomal origins (Lin et al., 2018). Diverse well-assembled reference genomes would also provide a fundamental tool for functional genomics, genetic engineering, and molecular breeding in this economically important genus (Zhang et al., 2019). It would also improve phylogenomic analyses of the *Populus* pan-genome (Pinosio et al., 2016; Zhang et al., 2019), without the need for reliance on reference-guided mapping and variant calling based solely on the *P. trichocarpa* reference. Recent advances in approaches to whole genome sequencing, including chromosome conformation capture (Hi-C: van Berkum et al., 2010) and long-read sequencing offer a means to go beyond fragmented draft genomes and generate nearly comprehensive *de novo* assemblies (El-Metwally et al., 2014; Mardis, 2013).

*Populus tomentosa*, also known as Chinese white poplar, is indigenous and widely distributed across large areas of China (Gao et al., 2019). Moreover, it is also the first tree species planted in large-scale artificial plantations in China. Like other white poplars, *P. tomentosa* has become a useful model for genetic research on trees (An et al., 2011; Chen et al., 2018; Wang et al., 2014), but at present no genome sequence is available and the origin, evolution and genetic architecture of the *P. tomentosa* genome are unclear. It has been proposed as another species in the *Leuce* section, but is now considered a hybrid of *Populus alba* and *Populus adenopoda* or a tri-hybrid of the previous two taxa with *Populus tremula* (Dickmann and Isebrands, 2001). In recent years, similar conclusions were reached using limited molecular markers (He, 2005; Wang et al., 2014). However, there is as yet a lack of strong genomic evidence to support this hypothesis.

Here, we present *de novo* assembles for *P. tomentosa* (clone GM15) by the combined application of PacBio, Illumina and Hi-C sequencing platforms. We herein provide two high quality assemblies for all chromosomes whose phylogenetic affinities demonstrate the hybrid origin of this species. From previous phylogenetic analysis on chloroplast genomes of seven *Leuce* poplars (Gao et al., 2019), we deduced that the ancestors of *P. tomentosa* are *P. adenopoda* (female parent) and *P*.*alba* var. pyramidalis (male parent). This contrasts with previous suggestions that the male parent is *P*.*alba* (Dickmann and Isebrands, 2001; Wang et al., 2014). Furthermore, we uncovered chromosome structural variations and allele-indels across *P. tomentosa* genome. These findings will help to elucidate the mechanisms of speciation in *Populus*, as well as the role of artificial and natural selection in poplar genome evolution. This study provides the first highly contiguous assembly of a *Populus* genome since publication of the original *P. trichocarpa* reference (Tuskan et al. 2006), expands our understanding of the unique biology of large woody perennials and provides a powerful tool for comparative biology, breeding and biotechnology.

## Results

### Genome Assembly, Anchoring and Annotation

To obtain a high-quality reference genome for *P. tomentosa* “LM50 (male clone),” we cultivated anthers (Li et al., 2013) and regenerated a plantlet (GM15) from which DNA was extracted for sequencing. We sequenced and assembled the genome through a combination of PacBio, Hi-C and Illumina methods. Using the PacBio platform (Chin et al., 2013), we constructed 20-Kb libraries to sequence the genome, generating 11 cells There were 6.24 million PacBio single-molecule reads (∼54 Gb), corresponding to ∼70 × coverage of the diploid genome, whose size was estimated to be ∼800 Mb by K-mer analysis (Supplementary Fig. S1 and Table S1). We then performed PacBio assembly using an overlap-layout-consensus method implemented in CANU (Koren et al., 2017; Wang et al., 2018a). The primary assembly was further refined using chromosome contact maps. Hi-C (*in vivo* fixation of chromosomes) technology (Dudchenko et al., 2017; Durand et al., 2016a; Putnam et al., 2016) was employed to construct chromosomes, then the assembly was corrected by genome data of *P. tomentosa* generated from an Illumina (HiSeq X-10) platform with Pilon over five iterations (Walker et al., 2014) (Table S2). A total of 38 chromosome-scale pseudomolecules were successfully constructed (Fig.S2, Table S3).

Simultaneously, transcripts derived from several white poplar species, including *P. alba, P. adenopoda, P. davidiana* (NCBI SRA), and *P. grandidentata* (NCBI SRA)—together with genomes derived from *P. alba* var. pyramidalis (Ma et al., 2019), *P. tremula* and *P. tremuloides* (Lin et al., 2018), and *P. trichocarpa* (Tuskan et al., 2006)--were used to assess the chromosome-scale pseudomolecules resulting from Hi-C analysis (based on co-phylogenetic analysis). The *P. tomentosa* genome was successfully separated into two subgenomes (2×19 choromosomes). Mapping of syntenic regions within the assembly showed clear chromosome-to-chromosome correspondence and also extensive synteny among different chromosomes, as expected for the highly duplicated *Populus* genome (Fig.1a).

**Figure 1.**
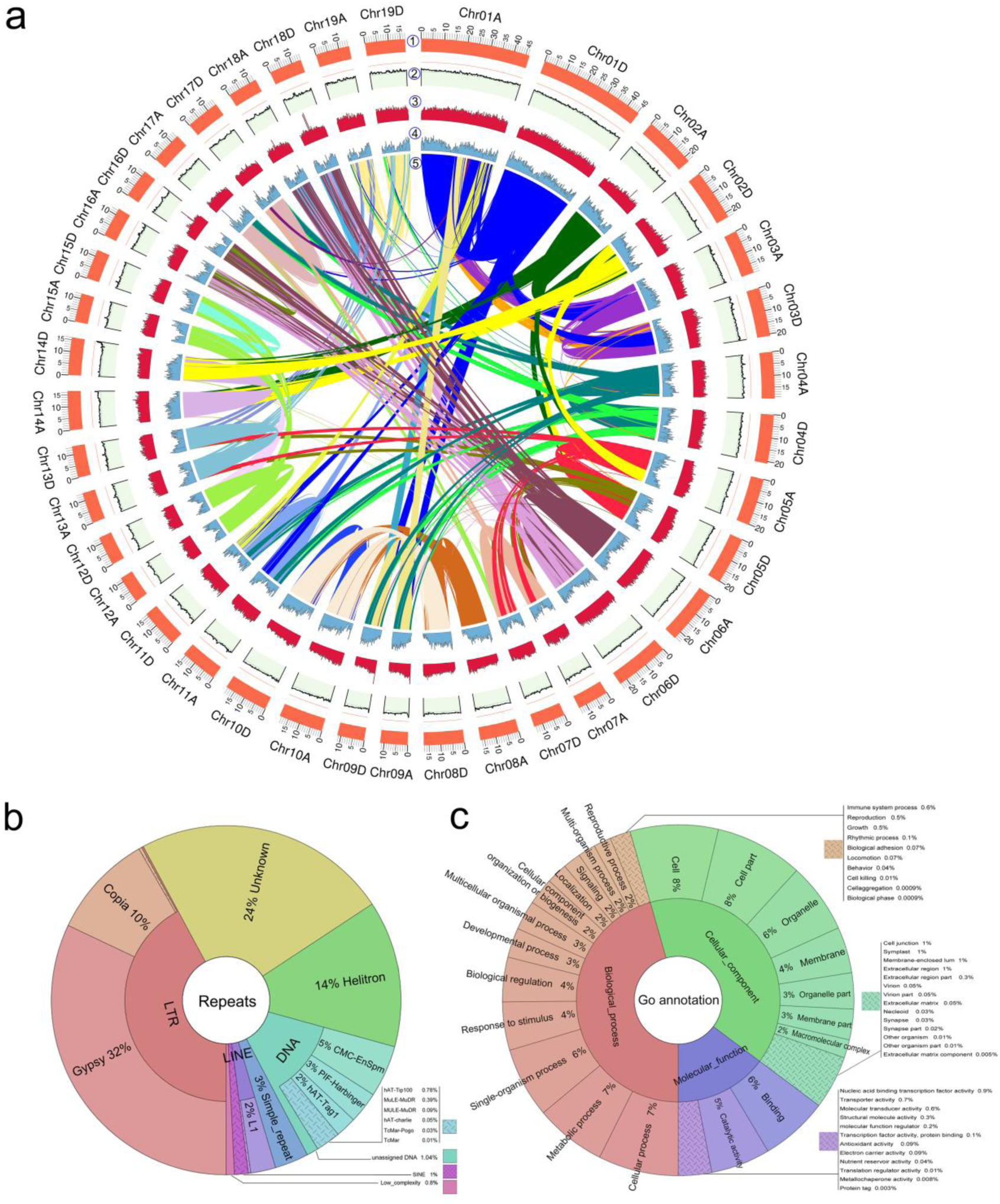
Characterization of the *Populus tomentosa* genome. **a**. *Populus tomentosa* genome overview. Genome features in 1-Mb intervals across the 38 chromosomes. Units on the circumference show megabase values and chromosomes. ① Choromosome karyotype; ② GC content (33.6 %. red line 50 %, green line 30 %). ③ Repeat coverage (45–1,937 repeats per Mb). ④ Gene density (3–164 genes per Mb). ⑤ The innermost parts are homologous blocks (1,463 genes) from paralogous synteny analysis. **b**. Distribution of repeat classes in the *P. tomentosa* genome. **c**. Distribution of predicted genes among different high-level Gene Ontology (GO) biological process terms.

Subsequently, we analyzed genome redundancy using the random best model of blasr (Chaisson and Tesler, 2012). The coverage depth distribution of BUSCO (Benchmarking Universal Single-Copy Orthologs) (Simao et al., 2015) for duplicated and single-copy core genes was identical, showing an expected Poisson distribution (Fig.S3). This indicates that these duplicated genes were not derived from assembling redundancy. More than 96% of complete genes identified by BUSCO could be detected in the genome. Meanwhile, the proportion of transcriptome data mapped to the genome reached 97.8%, suggesting that the genome was of high quality and nearly complete.

After integration of transcriptome, Illumina and Hi-C with the PacBio assemblies, a *de novo* assembly resulted in a draft diploid genome of 740.2 Mb for *P. tomentosa*. It was comprised of 38 chromosome-scale pseudomolecules that covered 92.1% of the 800-megabase genome of *P. tomentosa* (Table 1, Fig.1). Compared with previous poplar genome assemblies (Ma et al., 2019; Ma et al., 2013; Tuskan et al., 2006; Yang et al., 2017), the *P. tomentosa* assembly quality in the present study was improved significantly; the sizes of contig N50 and scaffold N50 reached 0.96 Mb and 18.91 Mb, with the longest contig and scaffold being 5.5 Mb, and 46.7 Mb, respectively (Supplementary Table S4).

**Table 1.**
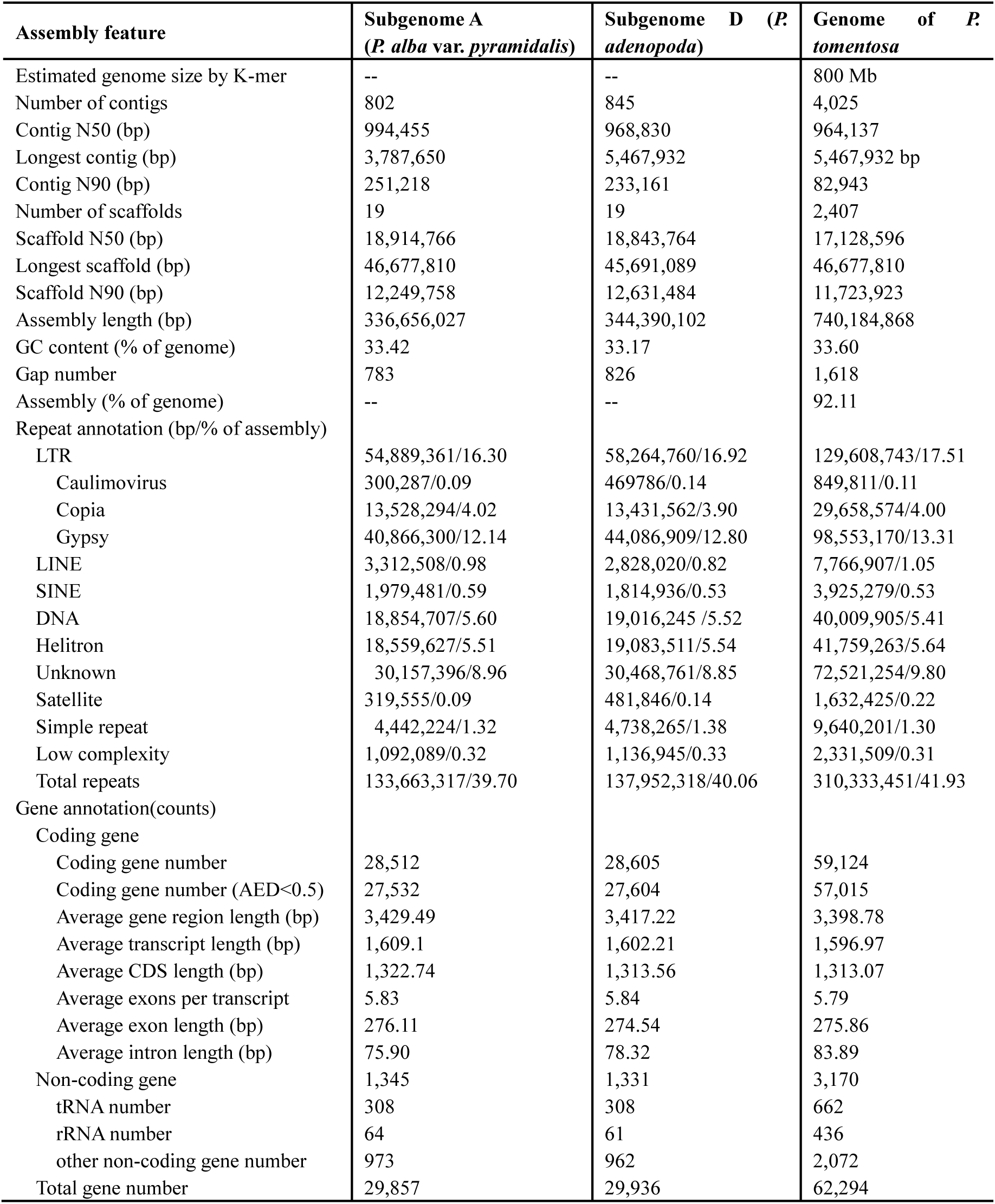
Statistics for the *P.tomentosa* draft genome.

| Assembly feature | Subgenome A<br>(*P. alba* var. *pyramidalis*) | Subgenome D (P.<br>*adenopoda*) | Genome of *P.*<br>*tomentosa* |
| --- | --- | --- | --- |
| Estimated genome size by K-mer | -- | -- | 800 Mb |
| Number of contigs | 802 | 845 | 4,025 |
| Contig N50 (bp) | 994,455 | 968,830 | 964,137 |
| Longest contig (bp) | 3,787,650 | 5,467,932 | 5,467,932 bp |
| Contig N90 (bp) | 251,218 | 233,161 | 82,943 |
| Number of scaffolds | 19 | 19 | 2,407 |
| Scaffold N50 (bp) | 18,914,766 | 18,843,764 | 17,128,596 |
| Longest scaffold (bp) | 46,677,810 | 45,691,089 | 46,677,810 |
| Scaffold N90 (bp) | 12,249,758 | 12,631,484 | 11,723,923 |
| Assembly length (bp) | 336,656,027 | 344,390,102 | 740,184,868 |
| GC content (% of genome) | 33.42 | 33.17 | 33.60 |
| Gap number | 783 | 826 | 1,618 |
| Assembly (% of genome) | -- | -- | 92.11 |
| Repeat annotation (bp/% of assembly) |  |  |  |
| LTR | 54,889,361/16.30 | 58,264,760/16.92 | 129,608,743/17.51 |
| Caulimovirus | 300,287/0.09 | 469786/0.14 | 849,811/0.11 |
| Copia | 13,528,294/4.02 | 13,431,562/3.90 | 29,658,574/4.00 |
| Gypsy | 40,866,300/12.14 | 44,086,909/12.80 | 98,553,170/13.31 |
| LINE | 3,312,508/0.98 | 2,828,020/0.82 | 7,766,907/1.05 |
| SINE | 1,979,481/0.59 | 1,814,936/0.53 | 3,925,279/0.53 |
| DNA | 18,854,707/5.60 | 19,016,245 /5.52 | 40,009,905/5.41 |
| Helitron | 18,559,627/5.51 | 19,083,511/5.54 | 41,759,263/5.64 |
| Unknown | 30,157,396/8.96 | 30,468,761/8.85 | 72,521,254/9.80 |
| Satellite | 319,555/0.09 | 481,846/0.14 | 1,632,425/0.22 |
| Simple repeat | 4,442,224/1.32 | 4,738,265/1.38 | 9,640,201/1.30 |
| Low complexity | 1,092,089/0.32 | 1,136,945/0.33 | 2,331,509/0.31 |
| Total repeats | 133,663,317/39.70 | 137,952,318/40.06 | 310,333,451/41.93 |
| Gene annotation(counts) |  |  |  |
| Coding gene |  |  |  |
| Coding gene number | 28,512 | 28,605 | 59,124 |
| Coding gene number (AED<0.5) | 27,532 | 27,604 | 57,015 |
| Average gene region length (bp) | 3,429.49 | 3,417.22 | 3,398.78 |
| Average transcript length (bp) | 1,609.1 | 1,602.21 | 1,596.97 |
| Average CDS length (bp) | 1,322.74 | 1,313.56 | 1,313.07 |
| Average exons per transcript | 5.83 | 5.84 | 5.79 |
| Average exon length (bp) | 276.11 | 274.54 | 275.86 |
| Average intron length (bp) | 75.90 | 78.32 | 83.89 |
| Non-coding gene | 1,345 | 1,331 | 3,170 |
| tRNA number | 308 | 308 | 662 |
| rRNA number | 64 | 61 | 436 |
| other non-coding gene number | 973 | 962 | 2,072 |
| Total gene number | 29,857 | 29,936 | 62,294 |

Using a combination of RepeatModeler (http://www.repeatmasker.org/RepeatModeler/) and RepeatMasker (http://www.repeatmasker.org/), 1,001,718 repeats were identified and masked (*de novo* identification, classification, and masking were run under default parameters, respectively). Collectively, these repeats were 307.6 Mb in size and comprised ∼41.5% of the genome (Fig.1a). Long-terminal repeats (LTR) were the most abundant, making up 17.5% of the genome. 13.3% of these were LTR/Gypsy elements, and 4.0% were LTR/Copia repeats. Second to LTR was unknown elements, making up 9.8% of the genome, followed by 5.6% Helitron repeats and 5.4% DNA elements (Fig.1b) (details in Supplementary Table S5).

To annotate genes, we trained the AUGUSTUS parameter model (Stanke et al., 2008) using single-copy core genes identified by BUSCO (Simao et al., 2015), followed by five rounds of optimization. We annotated the remaining unmasked *P. tomentosa* genome using a comprehensive strategy combining evidence-based techniques (RNA-Seq data and homologous protein) and *ab initio* gene prediction (Fig.1c). Using the MAKER2 gene annotation pipeline (Cantarel et al., 2008), we incorporated 73,919 protein sequences from two plant species and 137,918 transcripts assembled from *P. tomentosa* RNA-seq data. A total of 59,124 high quality gene models were identified, with an average coding-sequence length of 1.31 kb, 6.04 exons per gene, 430 amino acids (aa) per protein. There was 28.5% genome coverage with an average length of 210.8 Mb (Table 1, Supplementary Table S6, Table S7).

The annotated genes were associated with the three onotological classes: biological process, cellular components, and molecular functions (Fig. 1c). Using tRNAScan-SE (Lowe and Eddy, 1997) and RNAMMER (Lagesen et al., 2007), we predicted 662 tRNAs with a total length of 49,659 bp (average length per tRNA: 75 bp), and 436 rRNAs (106 28S rRNAs, 106 18S rRNAs, and 224 5S rRNAs) with a length of 610,293 bp. We also annotated 2,072 ncRNAs with a length of 218,117 bp using RfamScan (Kalvari et al., 2018). Finally, we performed alignments with protein databases using BLAT (Kent, 2002), with a maximum annotation ratio of 98.6% (Supplementary Table S8).

### Comparative Genomics and Evolution

We compared 19,594 gene family (59,124 genes) in the *P. tomentosa* genome with those of other three sequenced poplar genomes including *P. trichocarpa, P. euphratica*, and *P. pruinosa* using OrthoMCL (Zhang Z et al., 2003). A total of 22,386 gene families (142,738 genes) were identified by homolog clustering. In addition, 14,738 gene families (119,375 genes) were shared by all four poplar species, and 1,154 gene families consisting of 2,038 genes were found to be unique to *P. tomentosa* based on OrthoMCL “mutual optimization.” Similarly, 646/1,349, 179/261, and 399/1,041 gene families/genes were found to be unique to *P. trichocarpa, P. euphratica* and *P. pruinose*, respectively (Fig. 2a, Supplementary Table S9).

**Figure 2.**
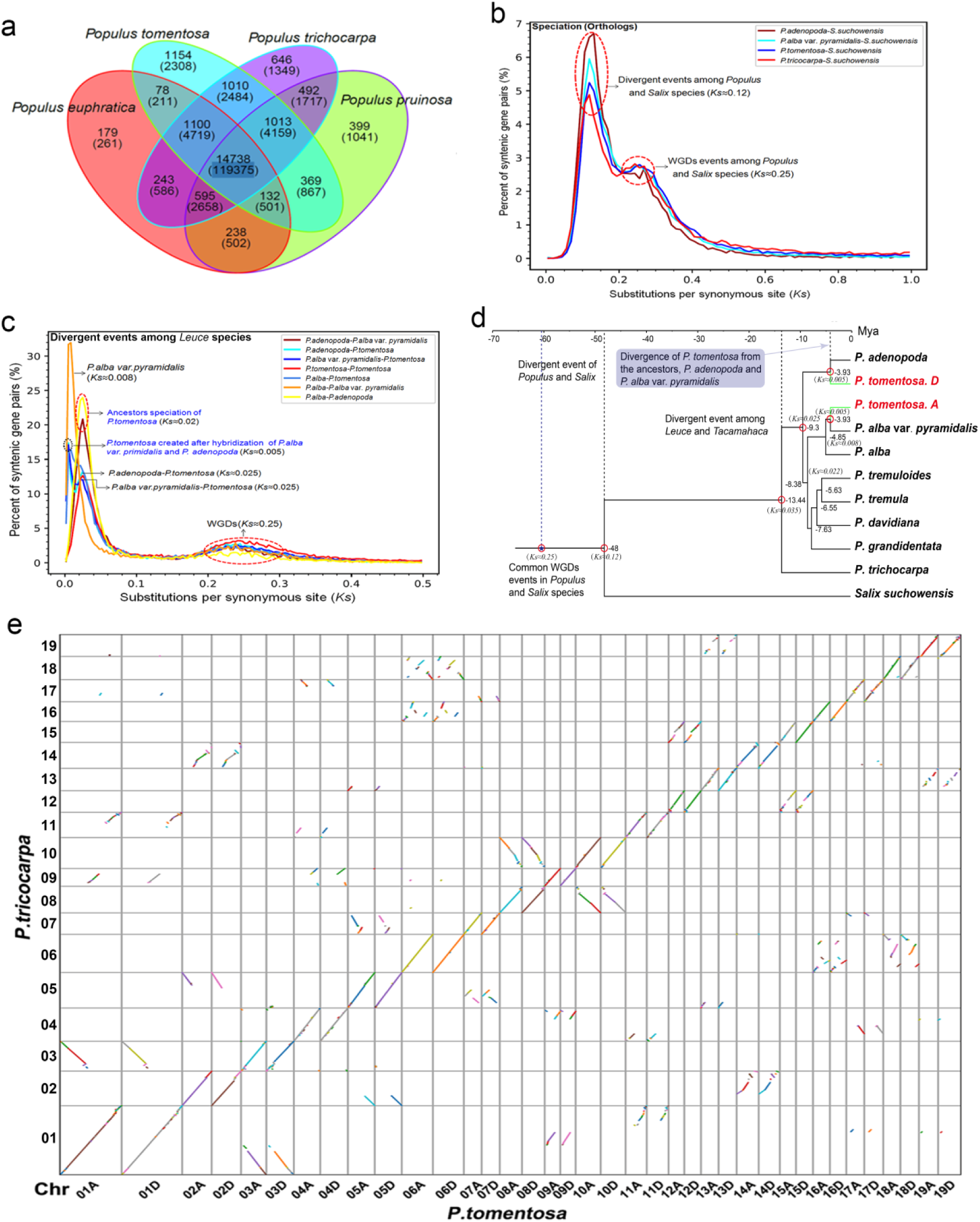
Comparative genomic analysis of *P. tomentosa* with other plants. (**a**) Shared gene families among *P. tomentosa* and three other poplars. The numbers indicate the number of genes (within each category). **b** Interspecific divergence in *Salicaceae* species, and intraspecific divergence in *Populus* species inferred by synonymous substitution rates (*Ks*) between collinear orthologous and paralogous pairs respectively. (**c**) Common genome duplication events (*Ks=* 0.25, ∼60 Mya) in *Salix* and *Populus* species (Dai et al., 2014; Ma et al., 2019), *P. tomentosa* speciation (*Ks* = 0.05, ∼3.93 Mya) and divergence events of other poplars as revealed through *Ks* analysis. (**d**) Inferred phylogenetic tree across 11 plant species using r8s (Sanderson, 2003). WGD events of *Salicaceae* species are placed. (**e**) Synteny between the *P. tomentosa* genome (the horizontal axis) and *P. trichocarpa* genome (the vertical axis). The *P. tomentosa* chromosomes were inferred to be syntenous with *P. trichocarpa* chromosomes based on orthologous genes from OrthoMCL analysis.

To address dates of divergence and duplication events in poplars, we conducted collinearity analysis of homologous gene pairs derived from *Populus* species vs. *Salix suchowensis* using MCScanX (Wang et al., 2012). From the *Ks* (synonymous substitution rate) distribution, we inferred a whole genome duplication event (based on paralogous pairs) and a species divergence event (based on orthologous pairs). The *Ks* distribution among syntenic genes of the four poplar species and *S. suchowensis* contained two peaks. One peak indicated that poplar and Salix species both underwent a common whole genome duplication (WGD) event (*Ks* ≈ 0.25), which was similar to a paleopolyploidization event occurred in the majority of flowering plants (Myburg et al., 2014; Otto, 2007). This result is consistent with a previous study on *Salix suchowensis* (Dai et al., 2014). Another peak represented the divergence between *Populus* and *Salix* occurred (*Ks* ≈ 0.12) (Fig.2b). Further analysis showed that section *Leuce* and *P. trichocarpa* have a divergence at *Ks* ≈ 0.035, and *P. adenopoda* and *P. alba* at *Ks* ≈ 0.025. Subsequently, *P. alba* var. pyramidalis is separated from *P. alba* at *Ks* ≈ 0.008. The hybridization event between *P. adenopoda* and *P. alba* var. pyramidalis subsequently occurred, followed by the emergence of *P. tomentosa* (*Ks* ≈ 0.005) (Fig. 2c).

To study the parental origin of *P. tomentosa*, we derived 1,052 single-copy orthologous genes from nine poplars, including *P. trichocarpa, P. alba, P. alba var. pyramidalis, P. adenopoda, P. tremula, P. tremuloides*, and *P. davidiana* for phylogenetic analysis. We constructed phylogenetic trees using *Salix suchowensis* as an outgroup using RaxML (Stamatakis, 2014), based on the GTR+GAMMA model (Allman et al., 2014) and maximum likelihood analysis (Guindon et al., 2010). Finally, referencing the fossil-based divergence time of *Populus* and *Salix* at 48 Mya (Boucher et al., 2003; Dai et al., 2014; Manchester et al., 1986; Manchester et al., 2006), we estimated dates for taxonomic divergence.

Phylogenetic analysis indicated that the divergence event between *Leuce* poplars and section *Tacamahaca* (*P. trichocarpa*) occurred at approximately 13.4 Mya. *P. adenopoda*, an ancestor of *P. tomentosa*, was the first to separate from the *Leuce* family as an independent clade approximately 9.3 Mya. Subsequently, the aspen tribe and white poplars tribe underwent a divergence event (approximately 8.4 Mya). Another ancestor of *P. tomentosa, P. alba var*. pyramidalis, gave rise to an independent variant of *P. alba* at approximately 4.8 Mya. Approximately 3.9 Mya, *P. tomentosa*, a new white poplar hybrid, was created by hybridization between *P. adenopoda* and *P. alba var*. pyramidalis (Fig. 2d).

We re-constructed phylogenetic trees of subgenome A, subgenome D of *P. tomentosa* and other poplars (Fig.S4), as well as for each of the corresponding 19 pairs of chromosomes (Fig.S5). All of these analyses supported the hypothesis that *P. tomentosa* genome originated from hybridization between *P. adenopoda* and *P. alba* var. *pyramidalis*. Based on the fact that the *P. alba* var. *pyramidalis* is a male tree, combining with our previous phylogenetic analyses of chloroplast genomes from *Populus* section *Leuce* (Gao et al., 2019), which indicated that P.tomentosa is much closer to *P. adenopoda* than other *Leuce* poplars, and following the law of maternal inheritance in chloroplast, we deduce that *P. alba* var. *pyramidalis* and *P. adenopoda* respectively played male and female parent roles in the hybrid formation of *P. tomentosa* during evolution.

Whole-genome synteny analysis revealed pairs of *P. trichocarpa*-homologous regions shared between chromosomes corresponding to the two subgenomes of *P. tomentosa*. A dot plot (Fig. 2e) indicated that most of the common linear segments of homologous chromosomes were shared between *P. trichocarpa* subgenome A and subgenome D. The diagonal distribution (“/”) indicated orthologous collinear genes in *P. tomentosa* and *P. trihcocarpa*, and other dispersed distribution-blocks in the dot plot, suggested the collinearity of paralogous genes on non-homologous chromosomes between the two poplars (Fig. 2e). These findings show that both of the *P. tomentosa* sub-genomes are highly syntenic with each other and *P. trichocarpa*.

To investigate potential recombination events between the two sub-genomes, the distance of 5,345 single copy orthologus genes from *P. tomentosa, P. alba* var. *pyramidalis* and *P. adenopoda* were estimated by *Ks*. We conducted tests whether recombination occurs between two subgenomes of *P. tomentosa* based on *Ks* comparisons; results indicated that no recombination events occurred within 4,309 gene loci (80.62%), recombination events did occur in 38 gene loci (0.87%), and 998 gene loci did not meet either of above two hypotheses (Table S10, Fig.S6). This suggests that the two parental genomes may be largely still intact in *P. tomentosa*, at least with respect to genic composition. Based on the speculation that *P. tomentosa* is F1 hybrid, therefore, it is reasonable to maintain basic independence and stability of the two sub-genomes in *P. tomentosa* genome.

### Chromosome structural variation in the *P. tomentosa* genome

To investigate the differences between subgenome A and subgenome D, we performed synteny analysis between paralogs in the *P. tomentosa* genome. This revealed collinear in-paralogous gene pairs, and suggested general collinearity at the sub-genome level, with dispersed collinear blocks among homologous and nonhomologous chromosomes (Fig. 3, center). We infer that these may have arisen from hybridization and duplication events occurred in *Populus* prior to their divergence.

**Figure 3.**
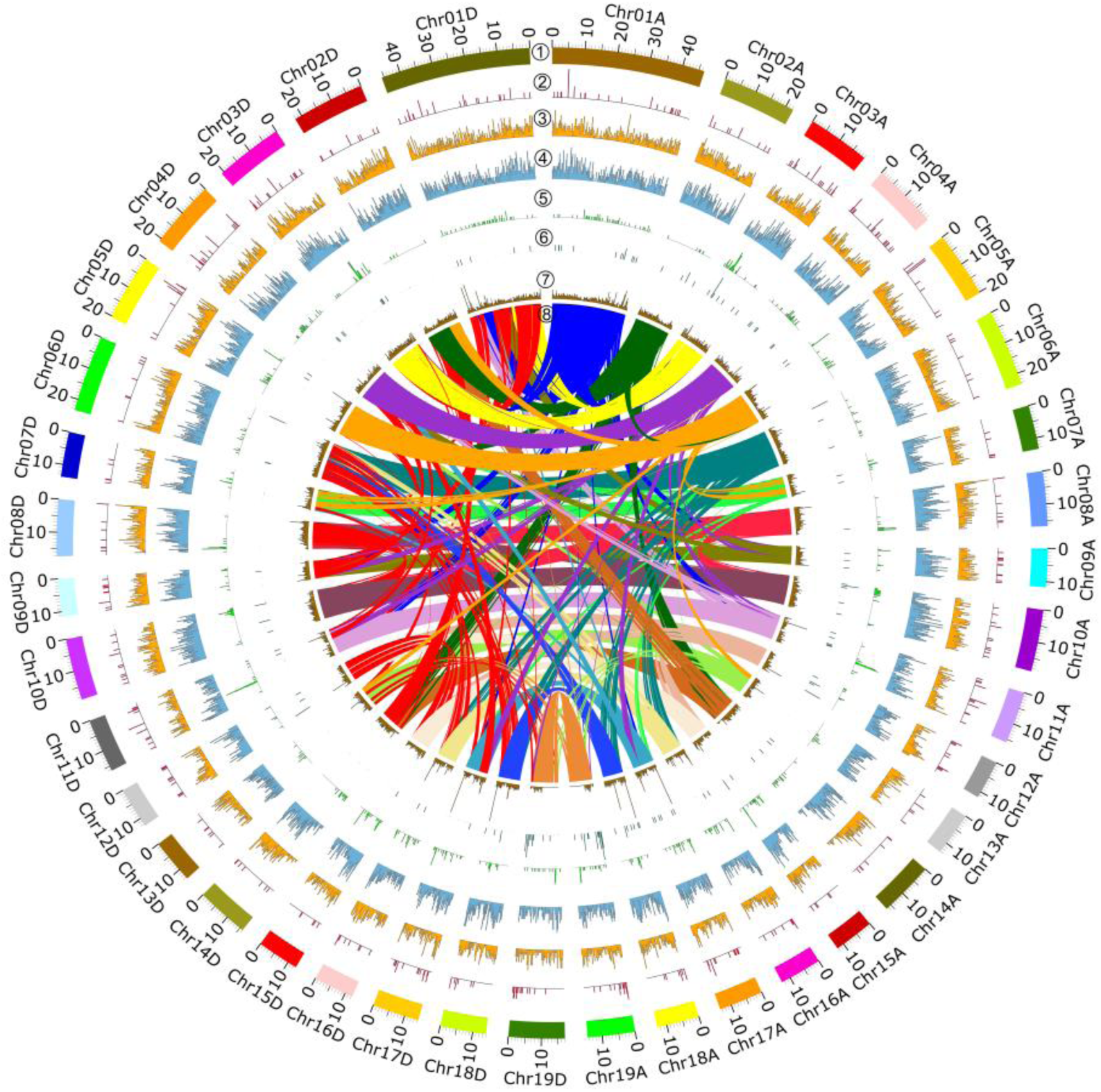
Synteny, structural variations and allele-indels analyses between subgenome A and subgenome D in *P. tomentosa*. ① Chromosome karyotype, ② Genomic distributions of copy number variations (CNV), ③ Genomic distributions of deletions (DEL), ④ Genomic distributions of insertions (INS), ⑤ Genomic distributions of inversions (INV), ⑥ Genomic distributions of translocations (TRANS), ⑦ Genomic distributions of indels between alleles of the two *P*.*tomentosa* subgenomes. ⑧ The inner part are synteny between subgenome A and subgenome D. The chromosomes of subgenome A were inferred to be syntenous with the chromosomes of subgenome D based on orthologous genes identified in OrthoMCL analysis.

To study genome-wide structural variation (SV), including copy number variation (CNV), deletions (DEL), insertions (INS), inversions (INV), and translocations (TRANS) among chromosome pairs (Fig. 3, rings 1-5), we conduted alignments using MUMmer, and subsequently called them out using SVMU (Structural Variants fromMUMmer) 0.3 (https://github.com/mahulchak/svmu). The results indicated that there were abundant chromosome structural variations in the *P. tomentosa* genome. Across the whole genome we detected 15,480 structural variations in total, of which INS (6,654) and DEL (6,231) accounted for the majority (83%). The other variant numbers were 1,602 and 694, and 299 for INV, TRANS and CNV, respectively, which together accounted for 27% of the total number of SVs observed (Table S11). The vast majority of INS, DEL, and CNV variations occurred between homologous chromosome pairs, whereas TRANS were generally seen between non-homologous pairs (Table S1, Fig.S7).

By plotting the distribution of five SV types along 38 *P. tomentosa* chromosomes, we noticed that total 299 CNVs presented irregular and sporadic distribution across the whole genome (Fig.3①). Relatively, there are more CNVs distributed on Chr01A and Chr01D, wherea fewer CNVs were distributed on Chr07A, Chr07D, Chr13A, Chr13D, Chr15A and Chr15D. We also noticed that most of DELs were almost evenly distributed through the whole genome, showing a slight preference for the telomere regions of Chr12A, Chr12D, Chr17A, Chr17D, Chr18A and Chr18D (Fig.3 ②). Similarly, INSs were present at high-density and showed a slight preference for telomere regions of Chr07A, Chr07D, Chr15A, Chr15D, Chr18A and Chr18D (Fig.3③). In contrast, INVs had a more uneven distribution across the genome (Fig.3 ④). INVs were more abundant on Chr01A and Chr01D, whereas their distribution was limited on other chromosomes. TRANS were very sparsely distributed on chromosomes, with only a few detected on Chr02D, Chr07D, Chr08D, Chr13D and Chr14D (Fig.3 ⑤).

Genome-wide scanning of indels (insertion/deletion) among allele paires was conducted using a web analytics tool (http://qb.cshl.edu/assemblytics/) (Nattestad and Schatz, 2016). A total 188,575 indels across 15,052 alleles were identified. We noticed that most of indels were evenly distributed on 38 chromosomes in *P. tomentosa*, and a high-density of indels were seen in regions close to telomere of Chr01A, Chr01D, Chr11A, Chr11D Chr12A, Chr12D Chr16A, Chr16D, Chr17A and Chr17D. Exceptions were the high-density indels seen on the middle and end of Chr18A and Chr18D (Fig.3 ⑥).

We performed GO enrichment analysis for total 15,480 SV variants using Plant GoSlim database, and detecd 23 GO categories significantly over-represented with respect to the whole set of *P. trichocarpa* genes (Fig.4). Ten of them (“motor activity”, “transporter activity”, “DNA binding”, “transport”, “metabolic process”, “lysosome”, “nuclear envelope”, “peroxisome”, “cell wall” and “extracellular region”) were over-represented in genes affected by INS, three (“chromatin binding”, “translation” and “ribosome”) were over-represented in genes affected by CNV, three (“hydrolase activity”, “response to biotic stimulus” and “lipid metabolic process”) were over-represented also in genes affected by both INS and TRANS, two (“cell differentiation” and “growth”) were over-represented also in genes affected by INV, two (“vacuole” and “circadian rhythm”) were over-represented also in genes affected by TRANS, one (“endosome”) was over-represented also in genes affected by both EDL and CNV, one (“carbohydrate binding”) was over-represented also in genes affected by EDL, CNV and TRANS, and one (“plasma membrane”) was over-represented also in genes affected by both CNV and TRANS. Overall, functional annotation showed enrichments associated with all of the major GO categories (“Molecular Function”, “Biological Process”, and “Cellular Component”).

**Figure 4.**
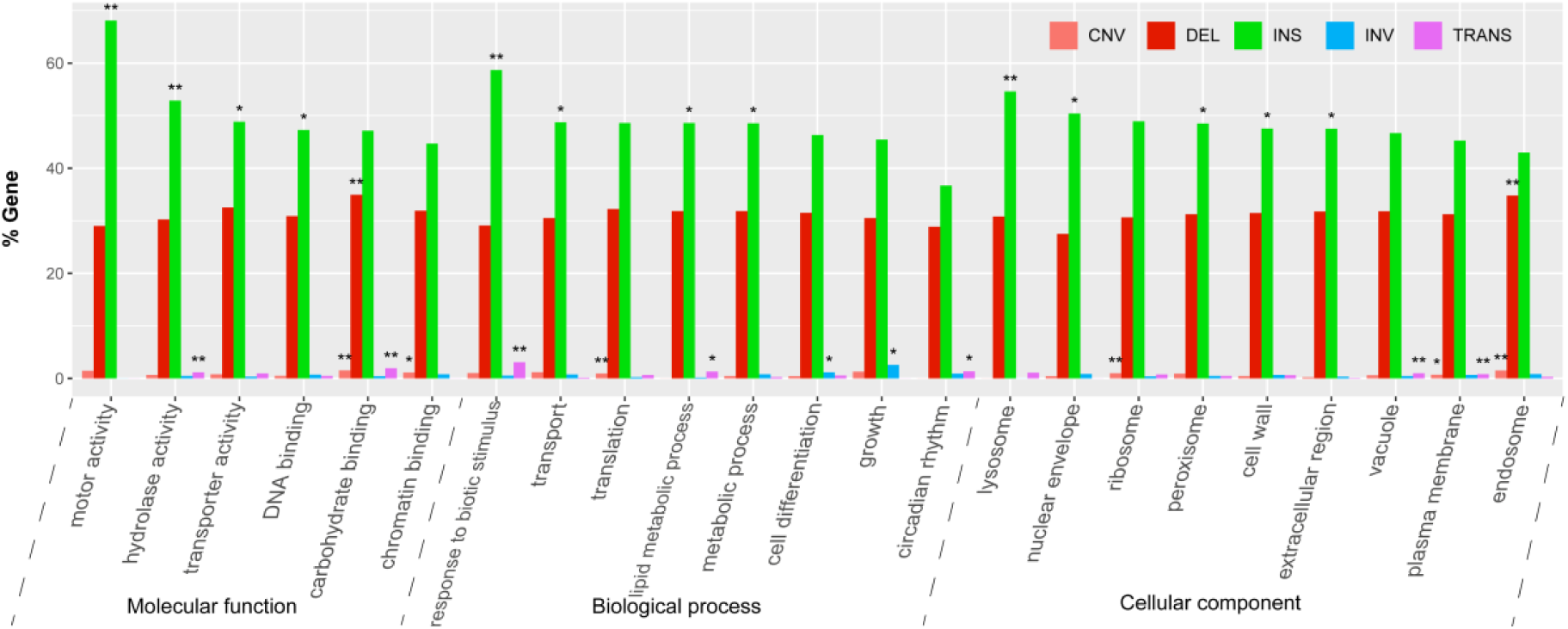
Functional classification of GO annotations of genes associated with chromosome structural variations. Frequencies of the Gene Ontology terms for which an over-representation has been observed when comparing the subsets of genes included in copy number variations (CNV), deletions (DEL), insertions (INS), inversions (INV), and translocations (TRANS) with respect to the complete dataset of *P. trichocarpa* annotated genes (ALL). * *P*-value<0.05, ** *P*-value<0.01.

## Discussion

*P. tomentosa*, also known as Chinese white poplar, is indigenous and widely distribueted and cultivated in a large area of China (Gao et al., 2019). Moreover, it was also the first tree species planted in large-scale artificial plantations in China. Here, we integrated advanced SMRT sequencing technology (PacBio), Illumina correction and chromosome conformation capture (Hi-C) to assemble a high quality genome. It contained 6.24 million PacBio single-molecule reads (∼54 Gb, ∼70 × coverage of whole genome) with a mean contig N50 of 0.99 Mb and a mean scaffold N50 of 18.91 Mb. The longest contig N50 was 5.47 Mb, and the longest scaffold N50 was 46.68 Mb. In comparion to several published poplar genomes, the assembly quality of *P. tomentosa* was better than those of *P. trichocarpa* (Tuskan et al., 2006), *P. euphratica* (Ma et al., 2013), *P. pruinosa* (Yang et al., 2017), *P. alba* var. pyramidalis (Ma et al., 2019). Even though the *P. alba* genome has a little bit longer contig N50 (1.18 Mb) than *P. tomentosa*, it has not been associated with specific chromosomes yet (Liu et al., 2019) (Supplementary Table 4). The whole genome size of *P. tomentosa* is 740.2 Mb, which is comprised of the sum of subgenome A (*P. alba* var. *pyramidalis*) and subgenome D (*P. adenopoda*). It obviously differs with those of *P. trichocarpa* (422.9 Mb), *P. euphratica* (497.0 Mb), and *P. pruinosa* (479.3 Mb), *P. alba* var. pyramidalis (464.0 Mb) and *P. alba* (416.0 Mb), which respectively consist of 19 chromosomes as they were resolved into a single haploid genome rather than two diploid subgenomes (Liu et al., 2019; Ma et al., 2019; Ma et al., 2013; Tuskan et al., 2006; Yang et al., 2017). However, this case is very similar to the genome of a hybrid poplar (84K) recently published, which was subdivided into two subgenomes (*P. alba* and *P. tremula* var. glandulosa) with a total genome size of 747.5 Mb (Qiu et al., 2019) (Table S3).

We presented evidence for divergence and duplication events in *Populus*, as well as within the *P. tomentosa* lineage. Like other many flowering plants (e.g., Myburg et al., 2014; Otto, 2007), *Salicaceae* species underwent a common palaeohexaploidy event, followed by a palaeotetraploidy event before the divergence of *Salix* and *Populus*. Subsequently, poplar speciation occurred gradually. *Leuce* poplars and *P. trichocarpa* differentiated from each other approximately 13.44 Mya (*Ks* ≈ 0.035). The ancestors of *P. tomentosa, P. adenopoda* and *P. alba var*. pyramidalis successively diverged from *Leuce* family approximately 9.3 Mya and 4.8 Mya. *Populus tomentosa* emerged from a hybridization event approximately 3.9 Mya. This finding differs from previous proposals on the origin of *P. tomentosa* (Dickmann and Isebrands, 2001; Wang et al., 2014).

Unlike other sequened poplars (Ma et al., 2013; Tuskan et al., 2006; Yang et al., 2017), the *P. tomentosa* genome consists of two parts, subgenome A (*P. alba* var. *pramidalis*) and subgenome D (*P. adenopoda*) (Fig. 1 and Table 1). Hi-C, as a chromosome conformation capture-based method, has become a mainstream techniqus for the study of the 3D organization of genomes (Belaghzal et al., 2017; Ma et al., 2018). Based on both Hi-C analysis (Fig. S2) and mapping of transcriptome (derived from *P. adenopoda* and genome data of *P. alba* var. *pramidalis)*, we were able to partion the *P. tomentosa* genome into two subgenomes. Phylogenetic analysis reveals the relationships among three white poplars (Fig. 2d, Fig. S4). Futher, 19 chromosome-phylogenetic trees provide solid evidence of the hybrid origin of the entire genome (Fig. S5). In addtion, the previous phylogenetic analysis of the chloroplast genome of *P. tomentosa* showed that it originated from *P. adenopoda* (Gao et al., 2019).

Our analysis of recombination events within genes showed that the *P. tomentosa* subgenomes have largely been maintained despite sharing the same nucleus for at least 3.93 million years. Comparision of 5,345 single copy orthologs from *P. tomentosa, P. alba* var. *pyramidalis* and *P. adenopoda* showed an absence of recombination among 99.13% of them (Fig. S6, Supplementary Table S10). This suggests that two subgenomes of *P. tomentosa* have been maintained relatively intact. Similarly, such karyotype stability has been observed in the paleo-allotetraploid *Cucurbita* genomes (Sun et al., 2017) and in newly synthesized allotetraploid wheat containing genome combinations analogous to natural tetraploids (Huakun et al., 2013; Zhang et al., 2013). However, this contrasts with the frequent homoeologous exchanges among subgenomes of allotetraploid cotton during its long evolutionary history after polyploidization (Li et al., 2015), as well as the considerable rearrangement events in many other polyploid plants (Simon and Wendel, 2014).

As in *Cucurbita* subgenomes (Sun et al., 2017), the karyotypic stability in *P. tomentosa* genome could be due to the rapid divergence between the two diploid genome donors in their repetitive DNA composition, which may have prevented meiotic pairing of homologous chromosomes and subsequent exchanges. This hypothesis may be supported by the observtion that dual-spindle formation in zygotes keeps parental genomes separate in some taxa but not others, such as for early mammalian embryos (Johansson et al., 2013; Reichmann et al., 2017). We also found that transposon abundance and distribution varied significantly. For example, LTR (Copia, Gypsy), LINE, SINE(tRNA), DNA(CMC-EnSpm, hAT-Ac, hAT-Tag1, PIF-Harbinger) and RC (Helitron).The few CNVs found in *P. tomentosa* may also be a reason for maintaining relative independence, stability and specifity of its subgenomes.

Transposable elements are also important factors that can cause CNVs, INSs and DELs due to their capacity to mobilize gene sequences within the genome (Kidwell and Lisch, 1997; Morgante et al., 2007; Pinosio et al., 2016), both in the wild and in breeding processes (Lisch, 2013; Olsen and Wendel, 2013; Sanseverino et al., 2015; Sun et al., 2018). Often these movements are associated with agronomically important traits such as skin or flesh color of the orange, grape, and peach (Butelli et al., 2012; Falchi et al., 2013; Kobayashi et al., 2004)—and their large variation among subgenomes may provide a degree of “fixed heterosis” that contributes to the high productivity and wide distribution of *P. tomentosa*.

## Methods

### Study system

The genus *Populus*, collectively known as the poplars, comprises six taxonomically distinct sections (*Leuce, Aigeiros, Tacamahaca, Leucoides, Turanga* and *Abaso*) consisting of nearly 30 tree species found in the Northern Hemisphere. Poplars often segregate into geographically and morphologically distinct subspecies and varieties due to their large natural range, and show considerable genetic variation in their adapative characteristics (Evans et al., 2014). As predominantly dioecious, wind-pollinated species with low barriers to crossability among consectional taxa, numerous natural and induced hybrids have been described (Dickmann and Isebrands, 2001). Poplars are pioneer species that have among the highest potential rate of biomass growth of temperate tree species (Lin et al., 2018). These traits are often enhanced in interspecific hybrids, promoting their commercial and ecological value, with multiple uses including biofuel, fiber, lumber, pulp, veneer, bioremediation, and windbreaks (Stettler et al., 1996). *Populus* generally exhibits rapid growth, easy propagation, a modest genome size (<500 Mb), and is amendable to genetic transformation (Tuskan et al., 2006). Moreover, *Populus* displays abundant genetic variation at different taxonomic levels: among sections within the genus, as well as among species, provenances, populations, individuals, and genes (Stettler et al., 1996). This diversity underlies local adaptation commonly found in the genus (Evans et al., 2014; Holliday et al., 2016; Wang et al., 2018b), and facilitates the study of species origins, evolution and divergence (Wang et al., 2016). These characteristics render *Populus* an ideal system for genetics and functional genomics studies of trees and other perennial plants (Ma et al., 2013).

*Populus tomentosa* Carr., an indigenous white poplar that is widespread in parts of China, is classified in section *Leuce* along with the aspens. It is a dominant species in many of the ecosystem it occurs in, and is widely distributed within a 1,000,000 km^2^ area in the Yellow River, Huaihe and Haihe regions (Gao et al., 2019). Its characteristics include rapid growth, a thick and straight trunk, environmental stress tolerance, and a long lifespan (typically 100-200 years, but sometimes over 500 years). These traits make *P. tomentosa* valuable from economic, ecological and evolutionary perspectives, with applications that include timber, pulp and paper, veneer, plywood, bioremediation, wind break, carbon capture, and prevention of soil erosion.

### In vitro culture of anthers and regeneration

Branches with floral buds were collected from an approximately 35-year-old *P. tomentosa* clone (LM50), which is a male elite individual in *P. tomentosa* tribe, on January 5, 2015, and cultured in containers with clean water at 24 ± 1 °C under 16/8 h light/dark conditions in a greenhouse. We referenced previous anther induced regenerated system (Li et al., 2013) and adjusted the culture medium. After determination of microspore developmental stages and pretreatment, anthers were cultured on callus induction medium (H medium, with 1.0 mg/L 6-BA, 1.0mg/L NAA, 5.5g/L agarose, and 30 g/L sucrose, pH 5.8) in the dark at 24 ± 1°C and 60 –65 % relative humidity for 6 months during which anthers were transferred monthly onto fresh media. Callus was transferred to shoot induction medium (MS medium, with 0.5 mg/L 6-BA, 0.05 mg/L NAA, 5.5g/L agarose, and 30 g/L sucrose, pH 5.8) to induce adventitious buds under conditions in similar to those described above. After 5 weeks of culture, shoots regenerated and 1.5-2.0 cm shoot sections were cut off, and transferred to the shoot rooting medium (1/2 MS medium with 0.3 mg/L IBA 0.3mg/L, 5.5g/L agarose, and 20g/L sucrose, pH 5.8). Plantlets were generated 30-40 days later.

### Plant material and DNA/RNA extraction

The plantlet GM15 generated by *in vitro* culture and regeneration system of anther from LM50, was selected for genome sequencing. Tissue cultured plantlets of GM15 were grown in the tissue culture room under natural light supplemented with artificial light (16 h light/8 h dark). Fresh leaves, stems and roots were harvested, immediately frozen in liquid nitrogen and stored at - 80 °C for genome and transcriptome sequencing. Young leaves collected from *P. alba, P. alba var*. primidalis, *P. davidiana*, and *P*.*a denopoda* were also forzen in liquid nitrogen and stored at - 80 °C for transcriptome sequencing.

Genomic DNA and total RNA were extracted using the Qiagen DNeasy Plant Mini Kit and the Qiagen RNeasy Plant Mini Kit, respectively, following the manufacturer’s instructions (Qiagen, Valencia, CA, USA). DNA and RNA quality were evaluated by agarose gel electrophoresis and their quantities determined using a NanoDrop spectrophotometer (Thermo Fisher Scientific, Waltham, MA, USA).

### Genomic and RNA-seq library construction and sequencing

A total of 2 μg of intact genomic DNA was fragmented and used to construct short-insert PCR-free libraries (300 and 500 bp) following the manufacturer’s protocol (Illumina Inc., San Diego, CA, USA). The DNA libraries were used in paired-end sequencing (∼265 million paired-end reads, ∼40 G bases, ∼50× coverage) on the Illumina HiSeq X-10 sequencer. For SMRT sequencing, genomic DNA was fragmented using g-Tubes (Covaris, Inc., Woburn, MA, USA) and 18–22 kb DNA fragments were further purified using AMPure magnetic beads (Beckman Coulter, Fullerton, CA, USA). DNA Template Prep Kit 4.0 V2 (Pacific Biosciences, USA) was used to constructed 20-kb library following the manufacturer’s protocol (Pacific Biosciences, Menlo Park, CA, USA). Finally, 50 μg of high-quality genomic DNA was used to generate 11 SMRT cells and then sequenced on the PacBio Sequel platform (Pacific Biosciences). To assist prediction and annotation of genes from the *P. tomentosa* genome assembly, 2 μg of total RNA from various tissues was used to construct RNA-seq library and sequence on the Illumina HiSeq X-10 platform following the protocol of NEBNext Ultra RNA Library Prep Kit for Illumina (New England Biolabs Ipswich, MA, USA).

### *De novo* genome assembly and estimation of genome size

A total of 6.24 M PacBio post-filtered reads were generated from the 11 SMRT cells, producing a total of ∼6.24 million reads, and ∼54 G bases (∼ 70× coverage) of single -molecule sequencing data. *De novo* assembly was conducted using an overlap-layout-consensus method in CANU (Koren et al., 2017). The draft assembly was polished with Arrow (https://github.com/PacificBiosciences/GenomicConsensus) to improve accuracies The size of contig N50 reached 1.8 Mb, yielding a final assembly with a total length of 740.18 Mb. We estimated genome size by k-mer distribution analysis with the program Jellyfish (k = 17) using Illumina short reads, and obtained an estimate of 803.6 Mb by using the Genome Characteristic Estimation (GCE) program (Liang et al., 2013). For additional details about k-mer distribution, see Supplementary Table S2.

### Hi-C library sequencing and scaffold anchoring

The Hi-C library was prepared using standard procedures described as follows. A total of 700 ng of high molecular-weight genomic DNA was cross-linked *in situ*, extracted, and then digested with a restriction enzyme. The sticky ends of the digested fragments were biotinylated, diluted, and randomly ligated to each other. After ligation, cross-links were reversed and the DNA was purified from protein. Purified DNA was treated to remove biotin that was not internal to the ligated fragments. Biotinylated DNA fragments were enriched and sheared to a fragment size of 300–500 bp again before preparing the sequencing library, which was sequenced on a HiSeq X-10 platform (Illumina). This yielded a total of 430 M reads, and 65 G bases (∼81× coverage).

The Hi-C reads were first compared to the above draft genome using Juicer (Durand et al., 2016b). To account for high duplication level, only the aligned sequences with map-quality score >40 were used to conduct Hi-C association chromosome assembly (Dudchenko et al., 2017). Visualization was carried out using Juicebox (Durand et al., 2016a). In Hi-C assembling, parameters were set to -m haploid -t 5,000 -s 2 -c 19. In another words, sequences above 5,000 bp in length were assembled, followed by two rounds of correction, and finally split into 19 chromosomes. Misjoined sequences were split in the process of correction, causing contigs to increase and N50 falls. For some obvious chromosomal splitting errors, we performed local optimization using the assembly method described above. Subsequently, the assembly was polished with Arrow over three iterations using PacBio reads and finally corrected using Illumina short reads with Pilon over five iterations.

### Transcriptome assembly and genome annotation

RNA-seq reads were preprocessed using Cutadapt to remove contaminating sequences from adaptors and sequences with low base quality. We employed a combination strategy of *de novo* and genome-guilded, consisting of 1) HiSat2 (Pryszcz and Gabaldon, 2016) + StringTie (Pertea et al., 2015), 2) HiSat2 (Pryszcz and Gabaldon, 2016) + Trinity (Grabherr et al., 2011) genome-guild mode, and 3) Trinity (Grabherr et al., 2011) *de novo* mode, to assemble the transcripts. Combining all of the transcripts and removing redundant sequences by CD-HIT (Fu et al., 2012) with 95 % identity and 95 % coverage, a total of 137,918 transcriptional sequences (Table 5) were yileded. These transcripts were used as EST evidence for subsequent gene annotation.

We annotated *de novo* repeats using the RepeatModeler (running in default parameter environment), subsequently masked repeat library using RepeatMasker pipeline (running in default parameter environment), and predicted genes using the MAKER2 annotation pipeline. Homologous protein evidence for the MAKER2 pipeline were provided in the form of 73,919 non-redundant protein sequences from *Arabidopsis* (TAIR10) and *P. trichocarpa* (JGI3.0). The single copy core genes identified by BUSCO (Simao et al., 2015) were used to train the AUGUSTUS (Stanke et al., 2008) parameter model, and five rounds of optimization were carried out. MAKER2 pipeline was carried out with combining *ab initio* prediction, EST sequence alignmenst and protein sequence alignmetns, and finally integrated these data with AED score calculated for quality control (Hoff et al., 2016). tRNA and rRNA were predicted using tRNAScan-SE (Lowe and Eddy, 1997) and RNAmmer (Lagesen et al., 2007), respectively. Other non-coding RNAs were annotated using RfamScan. Functional annotation was performed by aligning protein sequences with the protein database using BLAT (Kent, 2002) (identity >30 %, and the E < 1e -5).

### Molecular phylogenetic tree, whole-genome duplication and divergence events, subgenome recombination test

We firstly collected genome data of *P. tremula* (Lin et al., 2018), *P. tremuloides* (Lin et al., 2018), *P. trichocarpa* (Tuskan et al., 2006), and *Salix_suchowensis* (Dai et al., 2014), and *de novo* transcriptomes assembly of *P. davidiana*, and *P. grandidentata, P. adenopoda, P. alba* and *P. tomentosa* (Table S12). Then we performed gene family clustering using OrthoMCL (default parameters) on protein sequences, and conducted further collinearity analysis of homologous gene pairs derived from genome data of poplars and *Salix suchowensis* using MCScanX (Wang et al., 2012)

Based on collinear homologous gene pairs, including interspecific orthologs and intraspecific paralogs without tandem repeats, we aligned protein sequences using MUSCLE (Edgar, 2004), then used PAL2NAL to carry out codon alignment (Suyama et al., 2006). The YN model-based *Ka* and *Ks* calculation was performed using KaKs_Calculator (Zhang et al., 2006). The *Ks* distributions of the collinear homologous pairs (inparalogs and orthologs) were used to infer whole genome doubling (WGD) events and divergence events in species genome.

Finally, we referencing 1052 single copy orthologous genes derived from 10 plant species, constructed a molecular phylogenetic tree using RAxML (Stamatakis, 2014) based on the GAMMA+GTR model. Assuming the divergence time of *Populus* and *Salix*–48 Mya (Boucher et al., 2003; Dai et al., 2014; Manchester et al., 1986; Manchester et al., 2006) as fossil calibration, we estimated dates for WGD and divergent events of poplar species using r8s (Sanderson, 2003).

In additon to, we slected 5,345 single copy orthologous genes, which are collinear allele pairs between two subgenomes of *P. tomentosa* (PtA, PtD), and are homologus to those of *P. alba* var. pyramidalis (PA) and *P. adenopoda* (PD), to measure their distances through *Ks*, to investigate poptential recombinantion between homologus gene pairs based two hypothese: (1) if meet Ks (PD-PtD) < Ks (PD-PtA) and Ks (PA-PtA) < Ks(PA-PtD), then support the expectation of no recombination events between orthologous genes. (2) if meet Ks (PD-PtD) > Ks (PD-PtA) and Ks (PA-PtA) > Ks (PA-PtD), then support the expectation of recombination events between orthologous genes. If none of the above conditions are true, it may be false positive homology, gene loss, imbalance of evolution rate, etc., which shall not be taken as the judgment condition.

### Whole-genome synteny, chromosomal structure variations and indels in alleles

We conducted genome-wide synteny analysis between *P*.*tomentosa* and *P*.*trichocarpa*, and subgenome synteny analysis between subgenome A and subgenome D in *P*.*tomentosa* using MCScanX (Wang et al., 2012). Genome-wide structural variations (insertion, INS; deletion, DEL; inversion, INV; translocation, TRANS; copy number variation, CNV) between corresponding chromosome pairs in subgenomes were detected using MUMmer, and chromosome variation was identified using SVMU (Structural Variants from MUMmer) 0.3 (https://github.com/mahulchak/svmu). Besed on previous synety anlysis of homologous, alleles on homologous chromosomes were aligned using muscle, and indels were called out using a web tool (http://qb.cshl.edu/assemblytics/) (Nattestad and Schatz, 2016).

## Supporting information

Kmer analysis of Populus tomentosa (GM15, an individual derived from male clone LM50 using anther induction approach)

Hi-C Whole Genome of Populus tomentosa (GM15 )

The cover depth of duplicated core genes and single-copy core genes

The phylogentic tree of subgenome A and D and other poplars V5

The phylogenetic trees of 38 chromosomes1

Recobination test (Ks-diffence)

Chromosome structural vaiants count

Kmer statistics of Populus tomentosa genome (GM15)

Genome assembly version and statistics of Populus tomentosa (GM15)

Sequencing technology and assembly statistics comparisions of four poplars genomes.

Whole genome of Populus tomentosa (GM15)

Repeats statistics of Populus tomentosa (GM15)

RNA assembly of Populus tomentosa (GM15).

Coding genes statistics of Populus tomentosa (GM15)

Gene annotation of Populus tomentosa (GM15)

Venn figure data of Populus tomentosa (GM15)and other poplars

Cafe summary-tree of Populus tomentosa(GM15) and other species

Total chromosme structural variation statistics between two subgenomes

White poplars genome and transcriptome data sources

## Acknowledgements

Financial support was provided by the National Science and Technology Major Project of China (No. 2018ZX08021002-002-004), the National Natrual Science Foundation of China (No. 31870652, 31570661, 31170631), the National High Technology Research and Development Program (No. 2013AA102703) and the Major State Basic Research Development Program (No. 2012CB114505). We thank Professor Zhiyi Zhang for his forward-looking suggestions, also thank Shuhua Mu and Zewu An for figure editing.

## Author contributions

Xinmin An designed and managed the project. Rengang Zhang led the genome sequencing, assembly and analyses. Jianfeng Mao designed and led evolutionary analyses. Steven H. Strauss contributed to scientific analysis and interpretation, and edited manuscript. Stephen R. Keller participed in writing manuscript. Kai Gao and Y.L. created anther plant. Kai Gao, Zhong Chen, J.L., X.Y., X.Y.Y., J.Z., T.Y.Z., T.G., S.H., D.Y.M., W.K., B.G., S.W.L., and Nada prepared and collected all plant materials. J.W., B.Q.L., W.H.L., and Q.Z.Y. prepared RNA samples. P.R., M.X.Y., and L.X.J., preformed transcriptome assembly and anlysis. Xinmin An, Kai Gao, and Zhong Chen wrote the manuscript with input from other authors. All authors approved the manuscript before submission.

## Data availability

The raw reads generated in this study have been deposited in the NCBI Sequence Read Archive (https://www.ncbi.nlm.nih.gov/sra) under the BioProject accession PRJNA613008. The genome assembly and annotation of P. tomentosa has been deposited at DDBJ/ENA/GenBank under the accession JAAWWB000000000. The transcriptome assemblies have been deposited at DDBJ/EMBL/GenBank under the accessions GIKW00000000 (*P. grandidentata*), GILB00000000 (*P. davidiana*), GIKX00000000 (*P. adenopoda*) and GILC00000000 (*P. alba*).

## Additional information

Supplementary information accompanies this paper

