## Supplementary figures and images for "Hybrid origin of *Populus tomentosa* Carr. identified through genome sequencing and phylogenomic analysis"

### Fig.S1-K-mer analysis.pdf

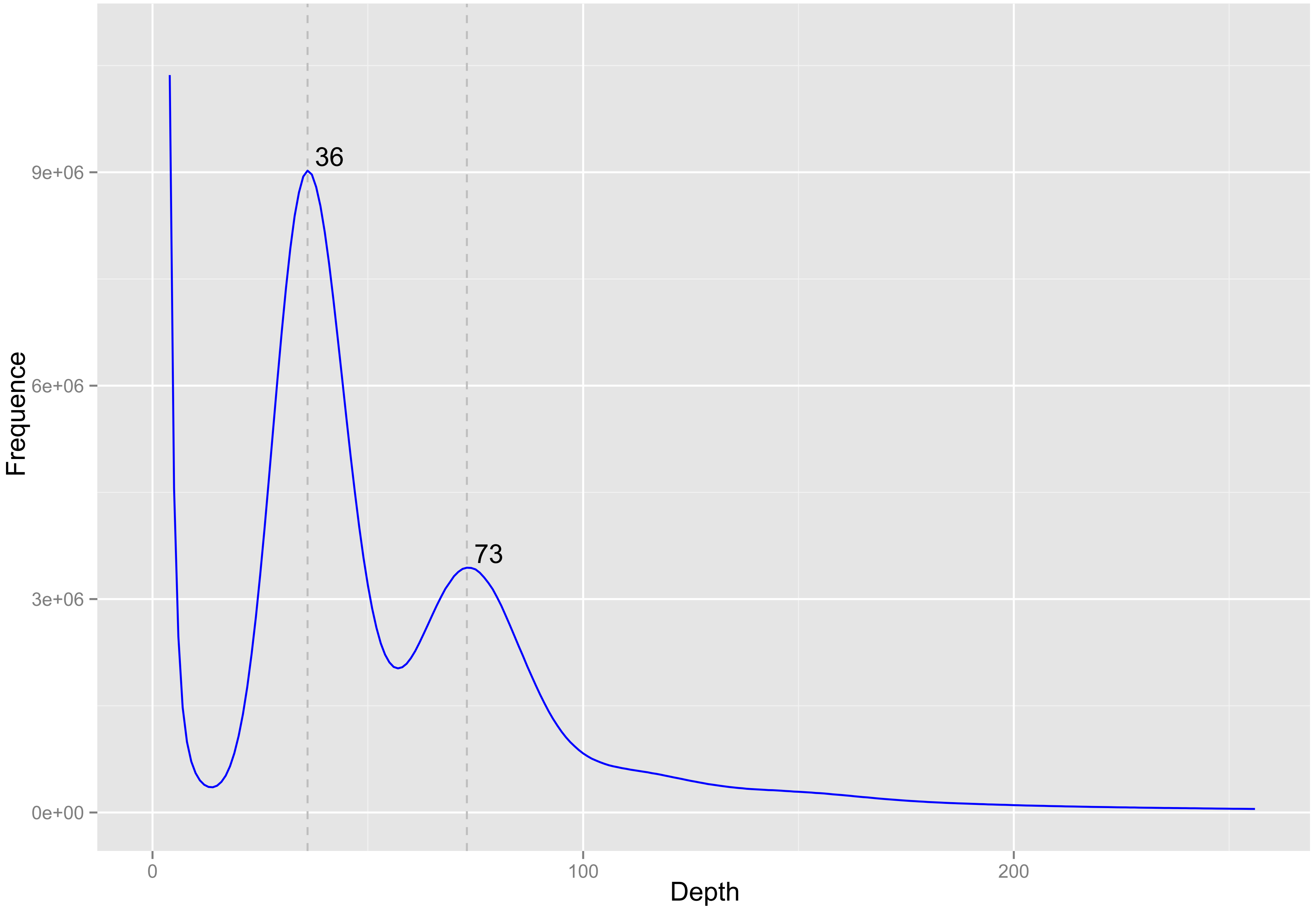

### Fig.S2-Hi-C Genome-WholeGenome-100K.pdf

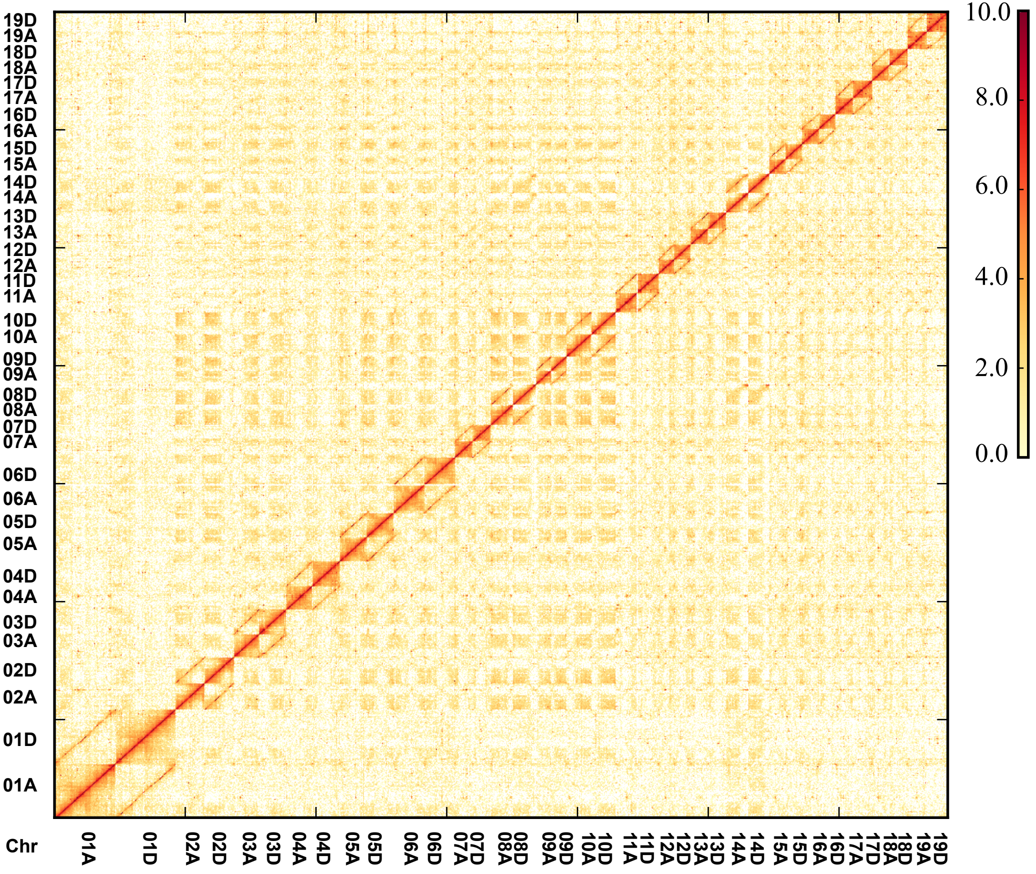

### Fig.S3-Poisson distribution.png

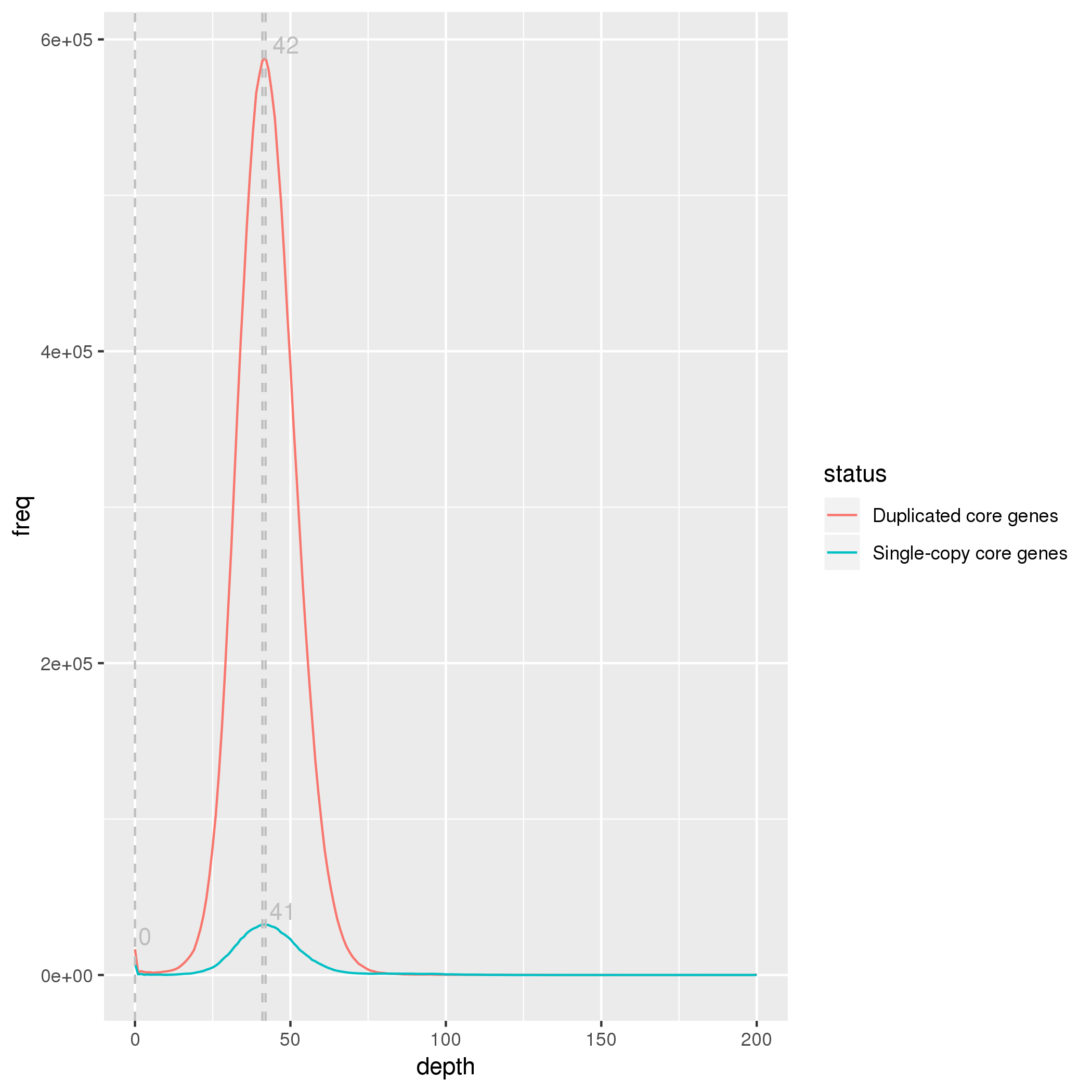

### Fig.S4 The phylogentic tree of subgenome A and D and other poplars V5.pdf

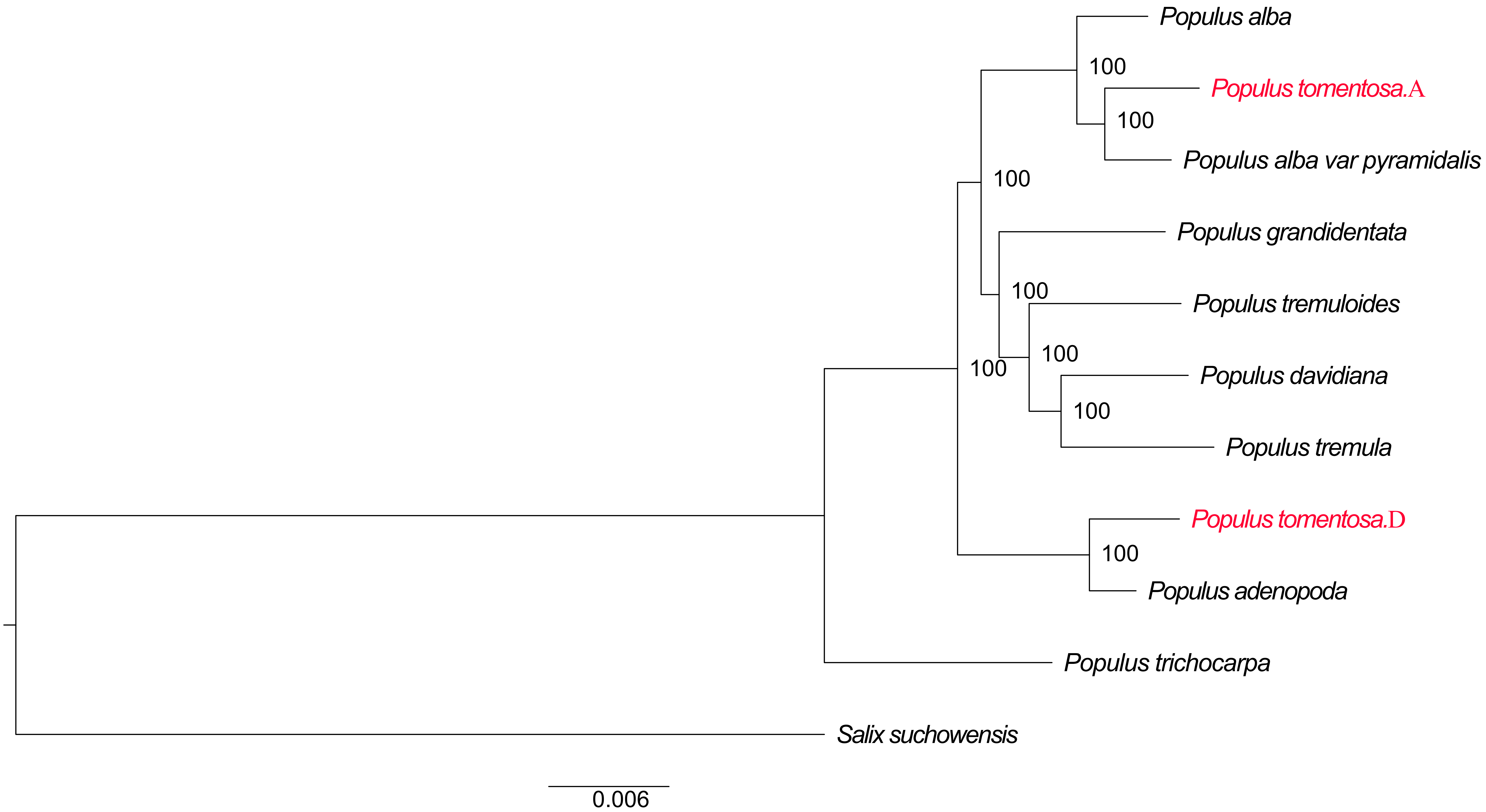

### Fig.S6 Recobination test (Ks-diffence).pdf

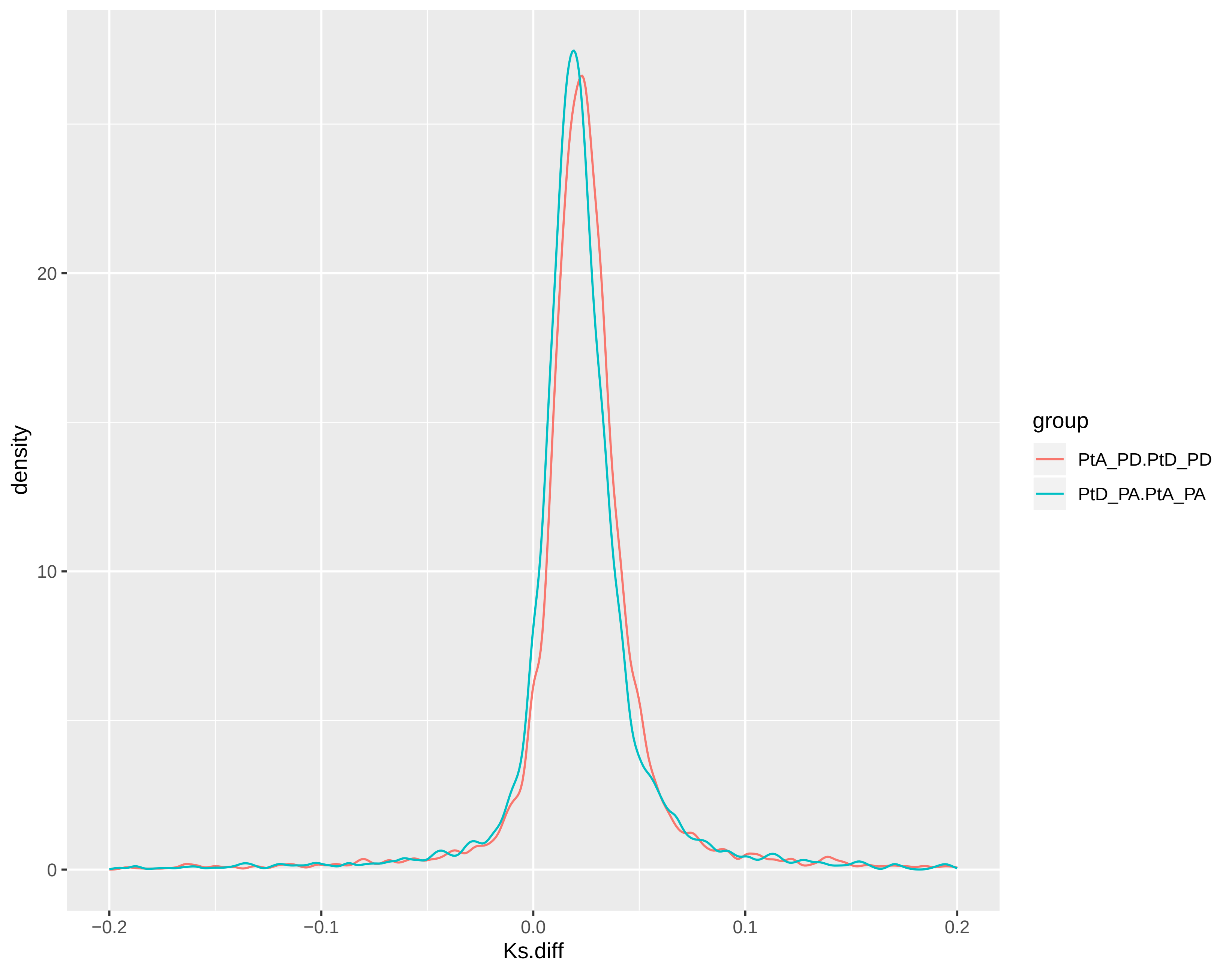

### Fig.S7 Chromosome structural vaiants count.pdf

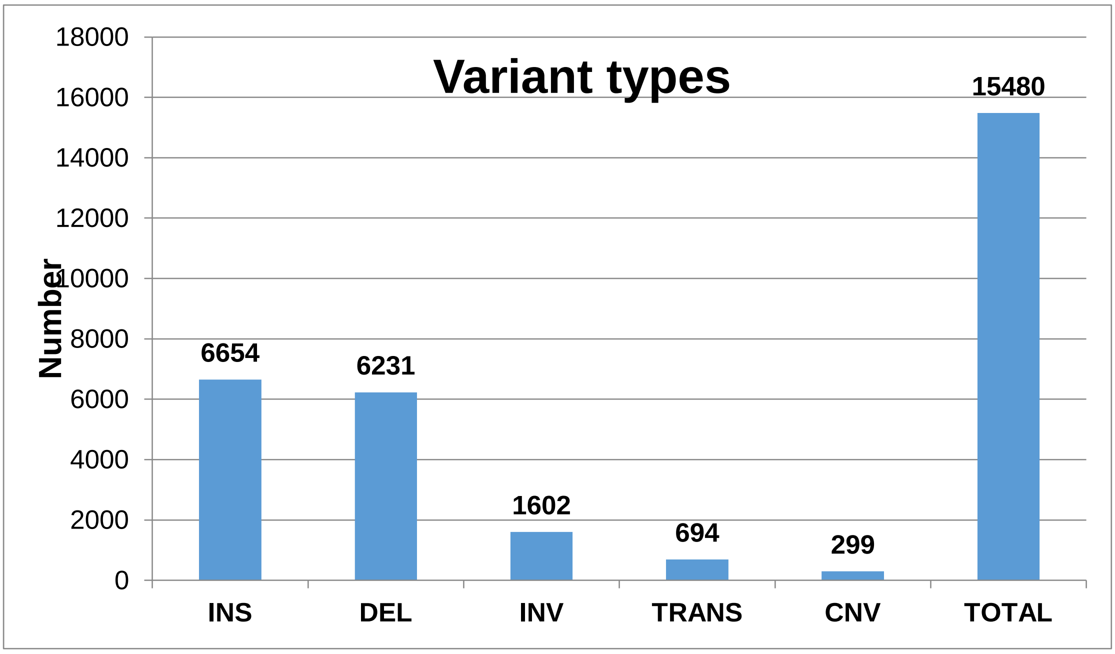
