## Supplementary material for "Hybrid origin of *Populus tomentosa* Carr. identified through genome sequencing and phylogenomic analysis": The phylogenetic trees of 38 chromosomes1: Fig.S5 The phylogenetic trees of 38 chromosomes1.pdf

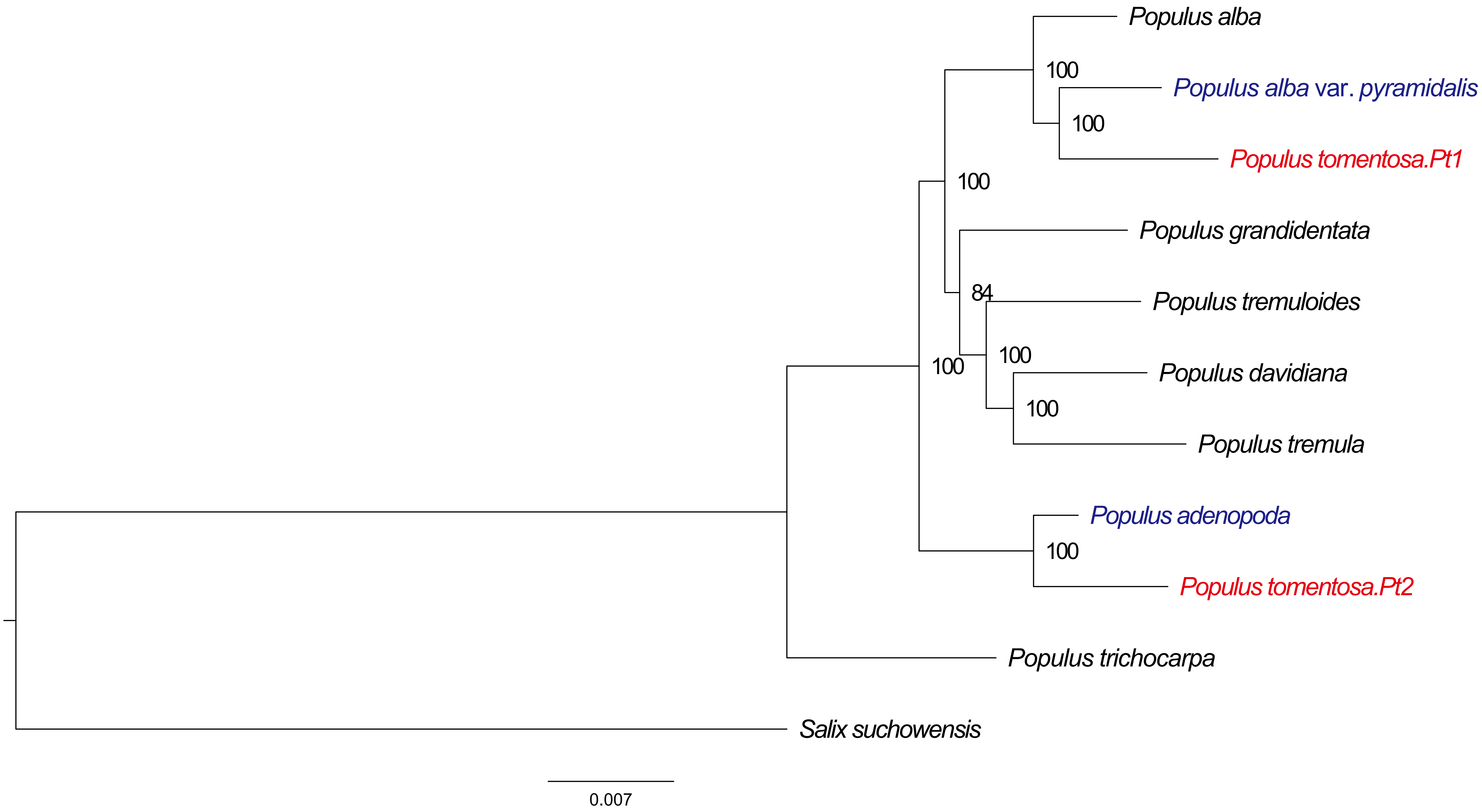
Populus alba
100
Populus alba var. pyramidalis
100
Populus tomentosa.Pt1
100
Populus grandidentata
84
Populus tremuloides
100
100
Populus davidiana
100
Populus tremula
Populus adenopoda
100
Populus tomentosa.Pt2
Populus trichocarpa
Salix suchowensis
0.007

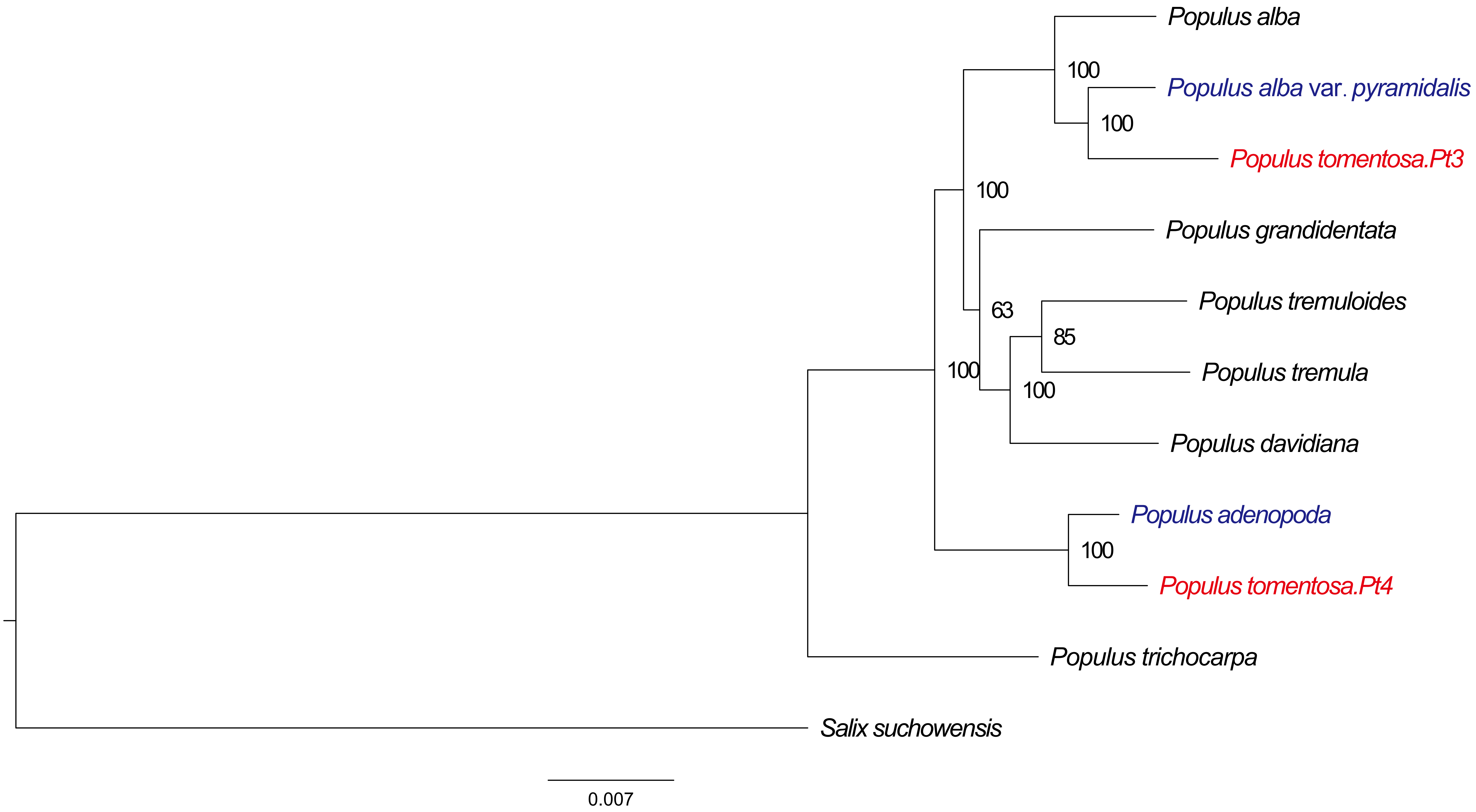

Populus alba
100
Populus alba var. pyramidalis
100
Populus tomentosa.Pt3
100
Populus grandidentata
63
Populus tremuloides
85
Populus tremula
100
100
Populus davidiana
Populus adenopoda
100
Populus tomentosa.Pt4
Populus trichocarpa
Salix suchowensis
0.007

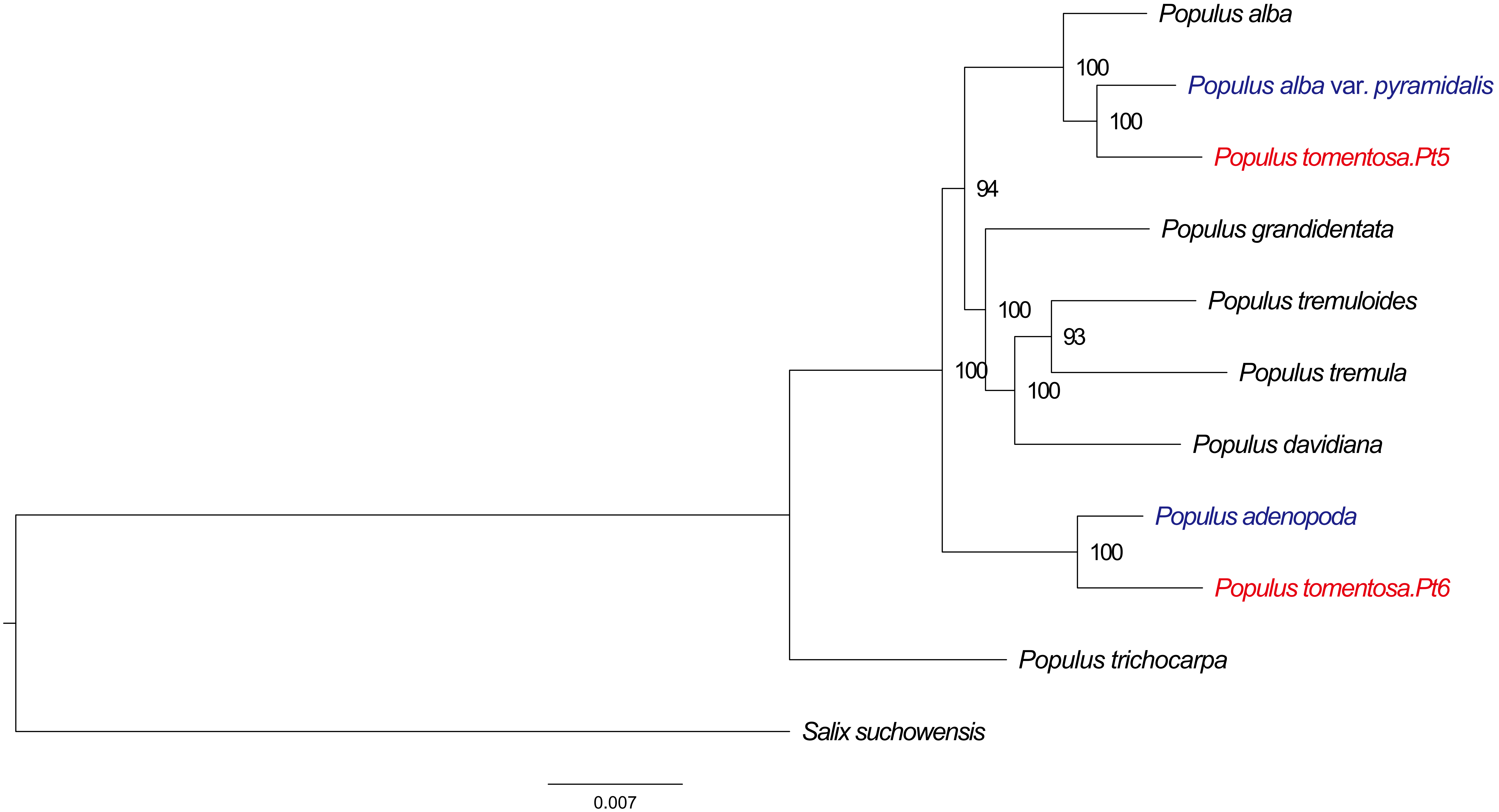
Populus alba
100
Populus alba var. pyramidalis
100
Populus tomentosa.Pt5
94
Populus grandidentata
Populus tremuloides
100
93
Populus tremula
100
100
Populus davidiana
Populus adenopoda
100
Populus tomentosa.Pt6
Populus trichocarpa
Salix suchowensis
0.007

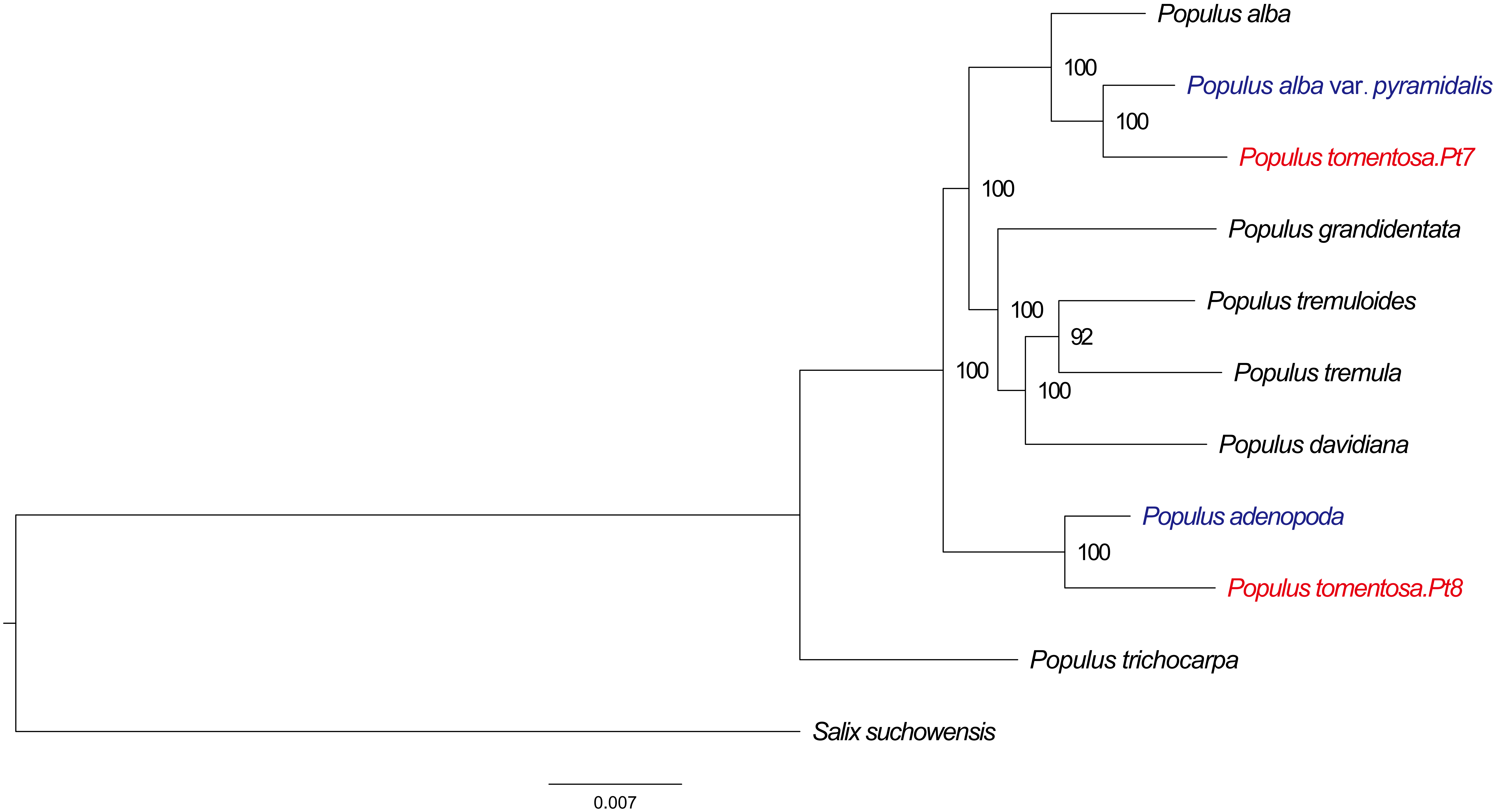

Populus alba
100
Populus alba var. pyramidalis
100
Populus tomentosa.Pt7
100
Populus grandidentata
100
Populus tremuloides
92
Populus tremula
100
100
Populus davidiana
Populus adenopoda
100
Populus tomentosa.Pt8
Populus trichocarpa
Salix suchowensis
0.007

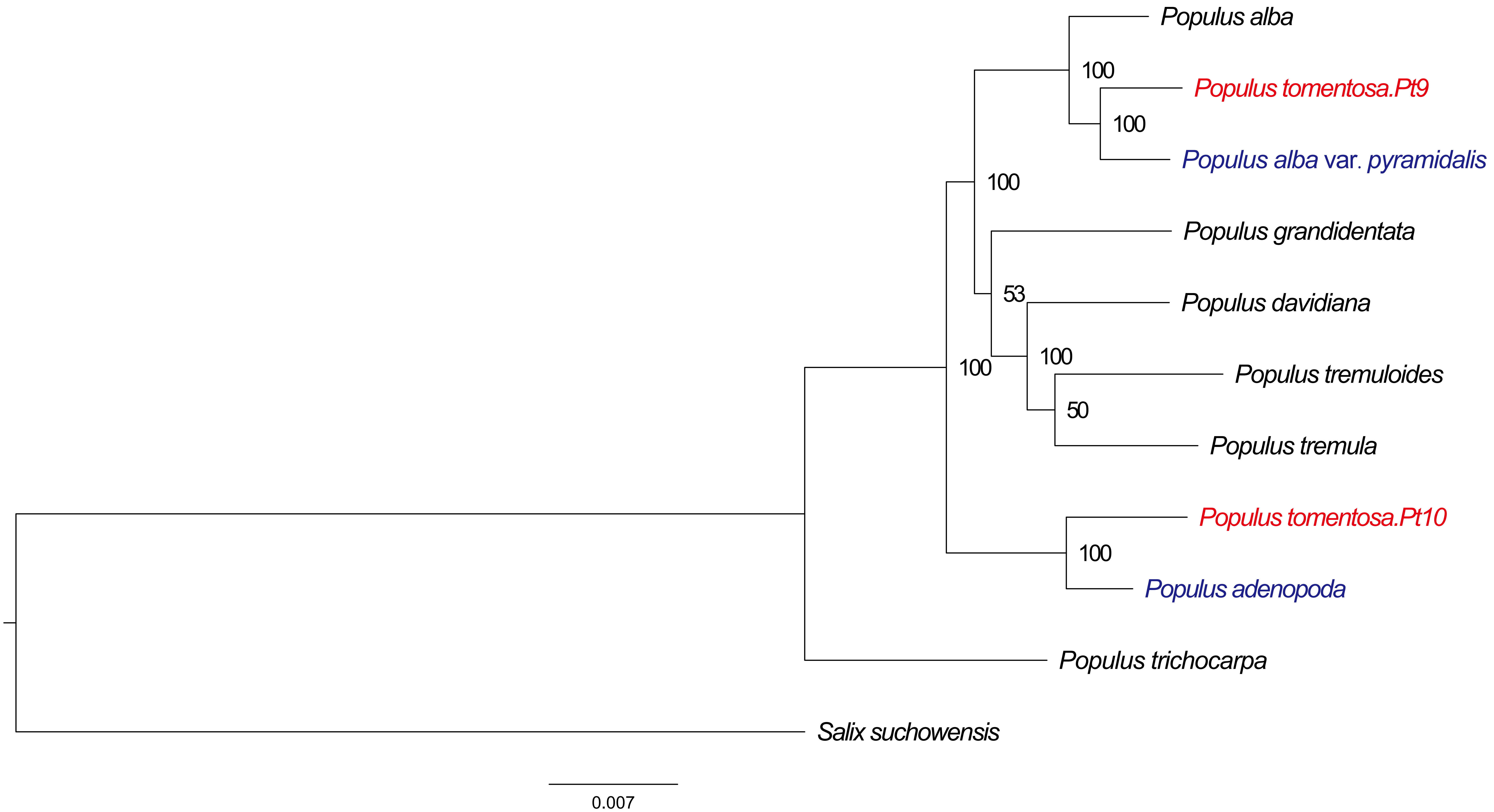
Populus alba
100
Populus tomentosa.Pt9
100
Populus alba var. pyramidalis
100
Populus grandidentata
53
Populus davidiana
100
100
Populus tremuloides
50
Populus tremula
Populus tomentosa.Pt10
100
Populus adenopoda
Populus trichocarpa
Salix suchowensis
0.007

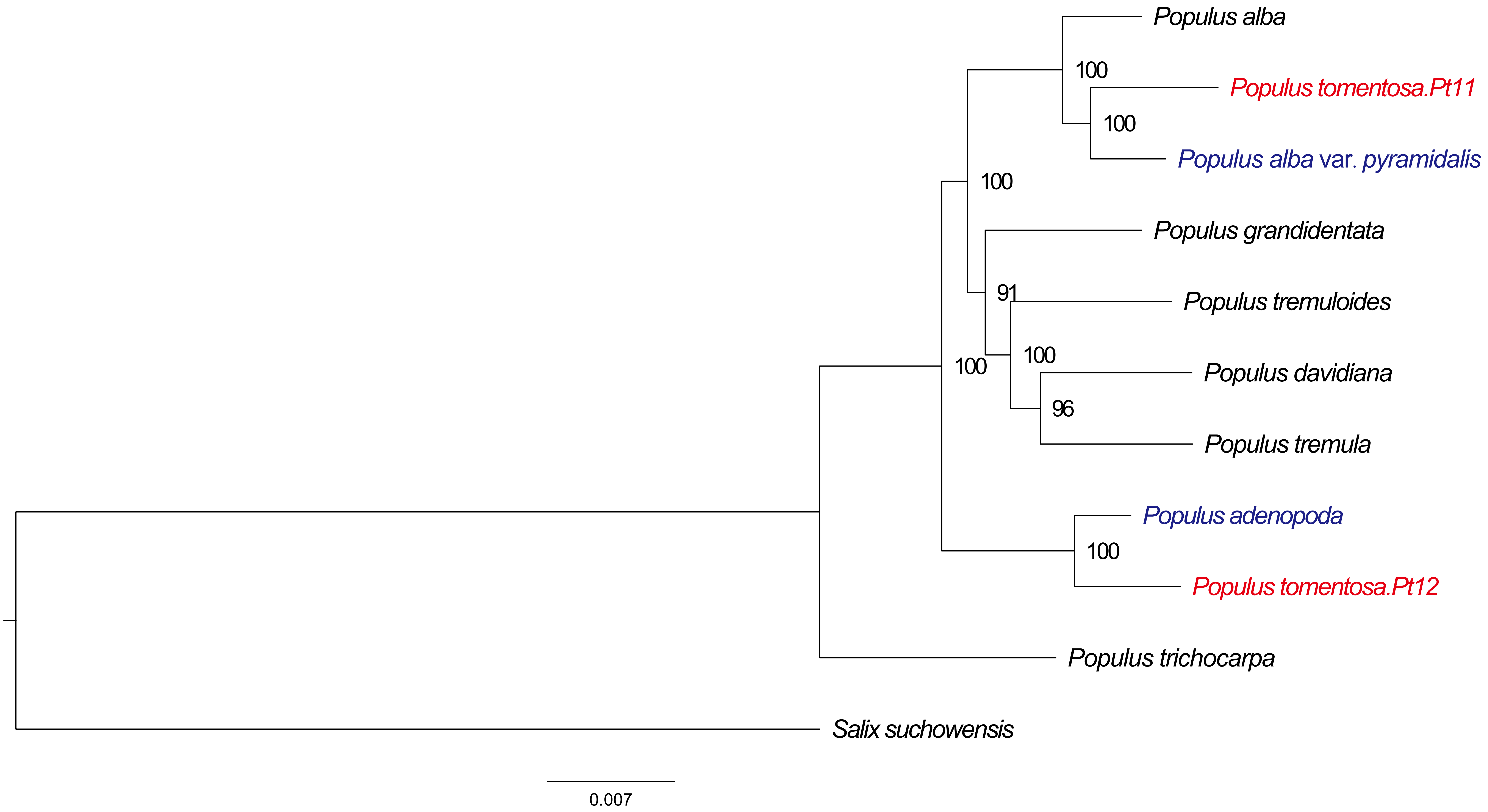
Populus alba
100
Populus tomentosa.Pt11
100
Populus alba var. pyramidalis
100
Populus grandidentata
91
Populus tremuloides
100
100
Populus davidiana
96
Populus tremula
Populus adenopoda
100
Populus tomentosa.Pt12
Populus trichocarpa
Salix suchowensis
0.007

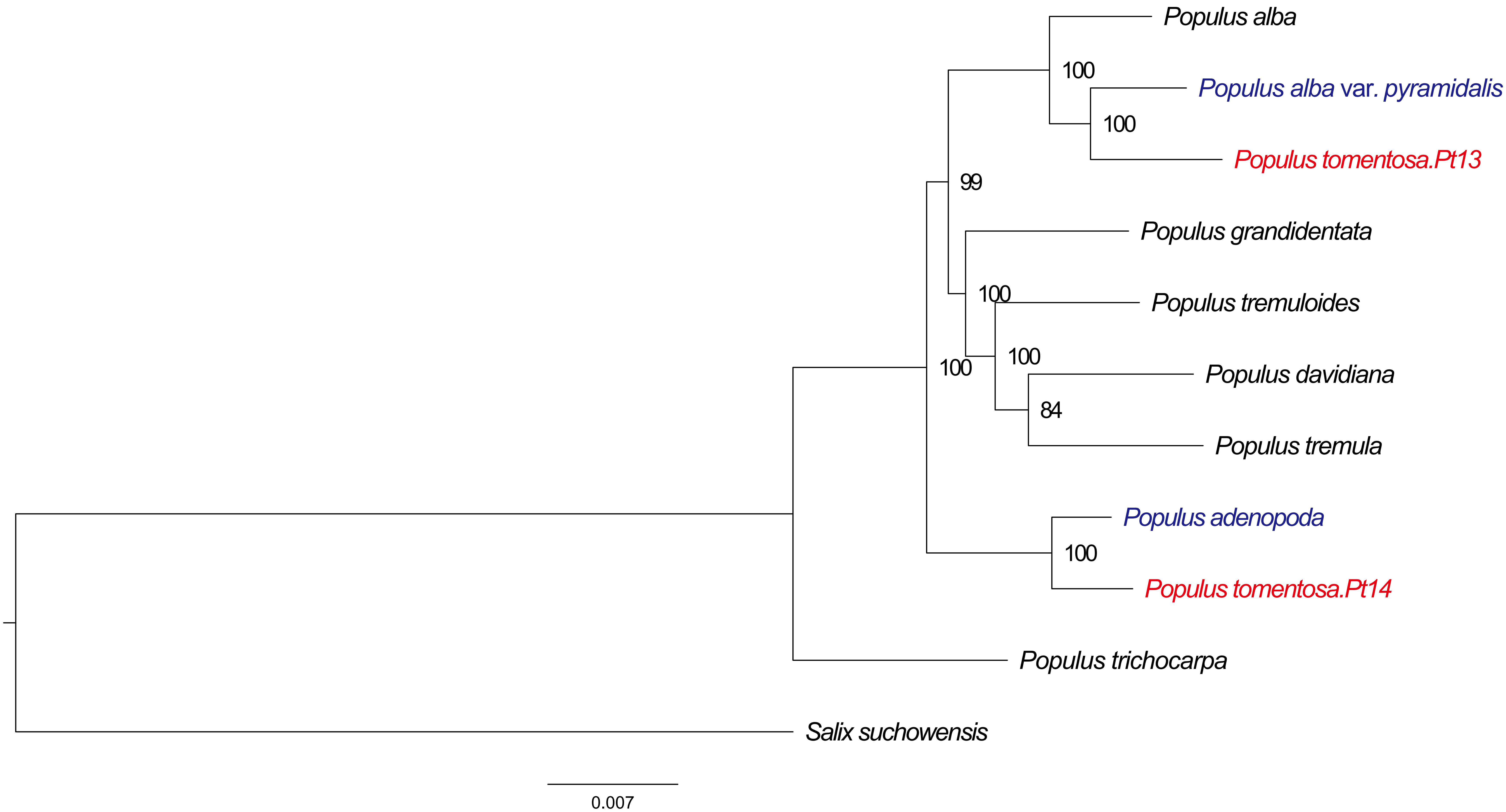

Populus alba
100
Populus alba var. pyramidalis
100
Populus tomentosa.Pt13
99
Populus grandidentata
100
Populus tremuloides
100
100
Populus davidiana
84
Populus tremula
Populus adenopoda
100
Populus tomentosa.Pt14
Populus trichocarpa
Salix suchowensis
0.007

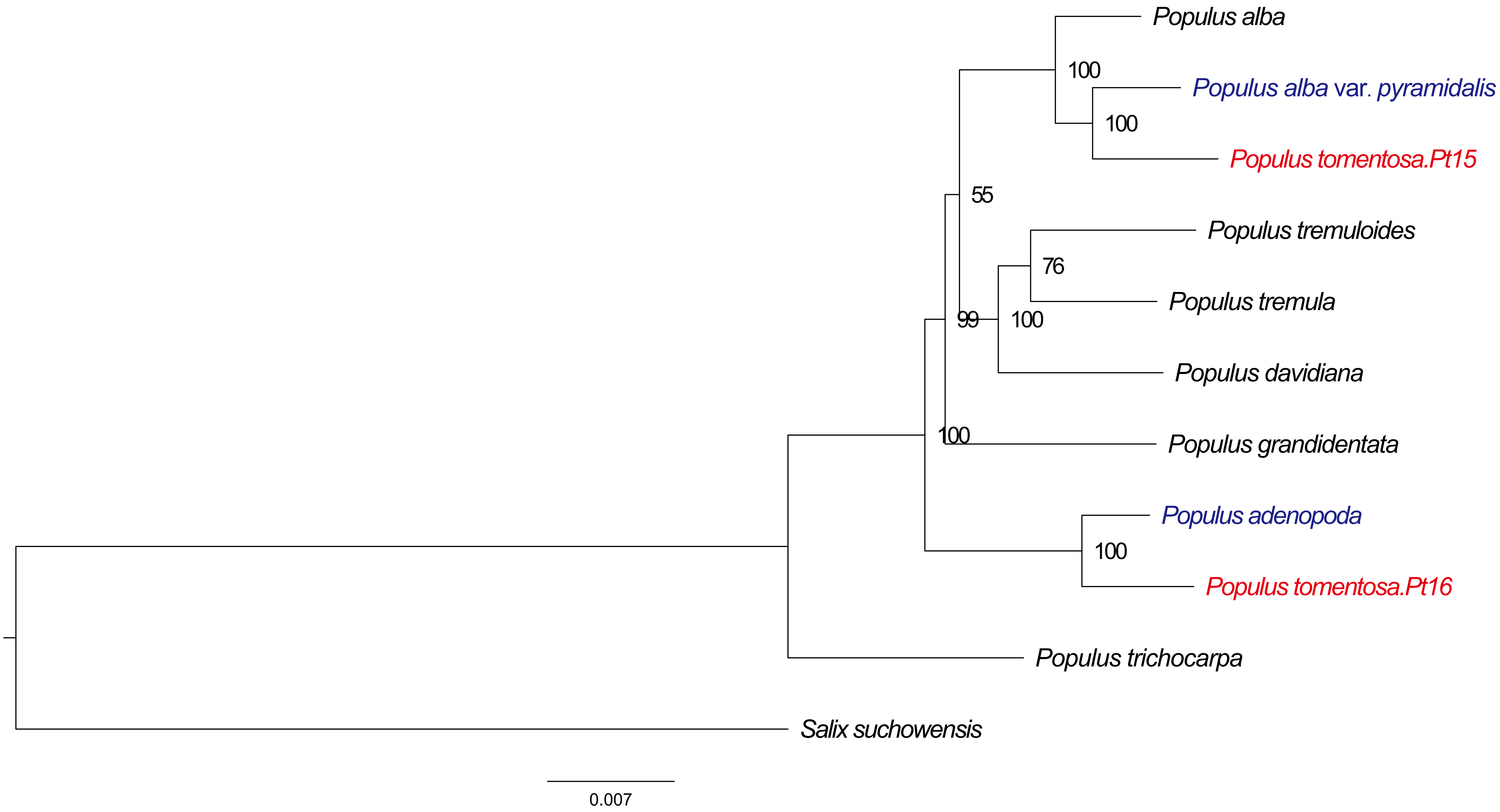
Populus alba
100
Populus alba var. pyramidalis
100
Populus tomentosa.Pt15
55
Populus tremuloides
76
Populus tremula
99
100
Populus davidiana
100
Populus grandidentata
Populus adenopoda
100
Populus tomentosa.Pt16
Populus trichocarpa
Salix suchowensis
0.007

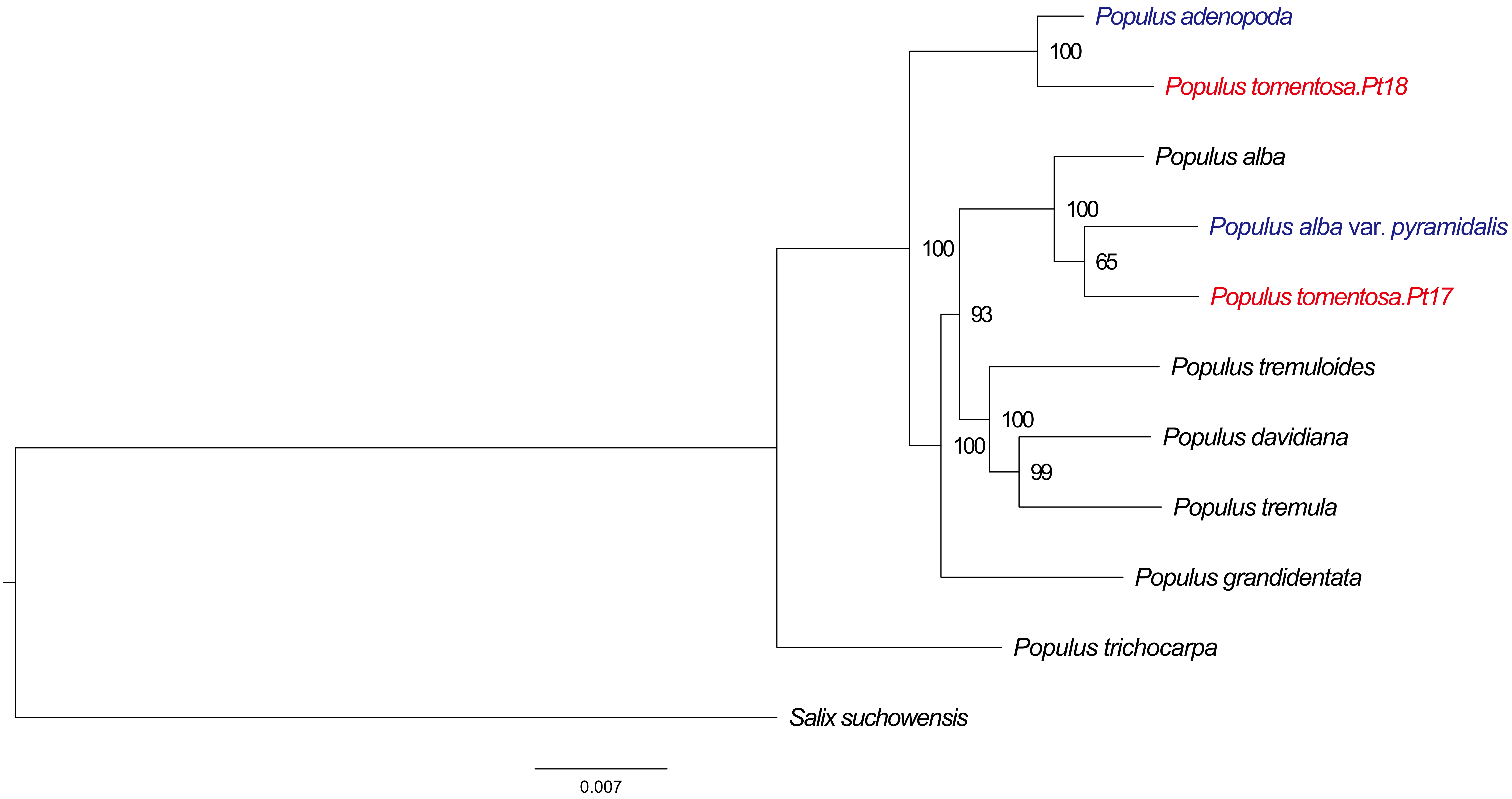

Populus adenopoda
100
Populus tomentosa.Pt18
Populus alba
100
Populus alba var. pyramidalis
100
65
Populus tomentosa.Pt17
93
Populus tremuloides
100
Populus davidiana
100
99
Populus tremula
Populus grandidentata
Populus trichocarpa
Salix suchowensis
0.007

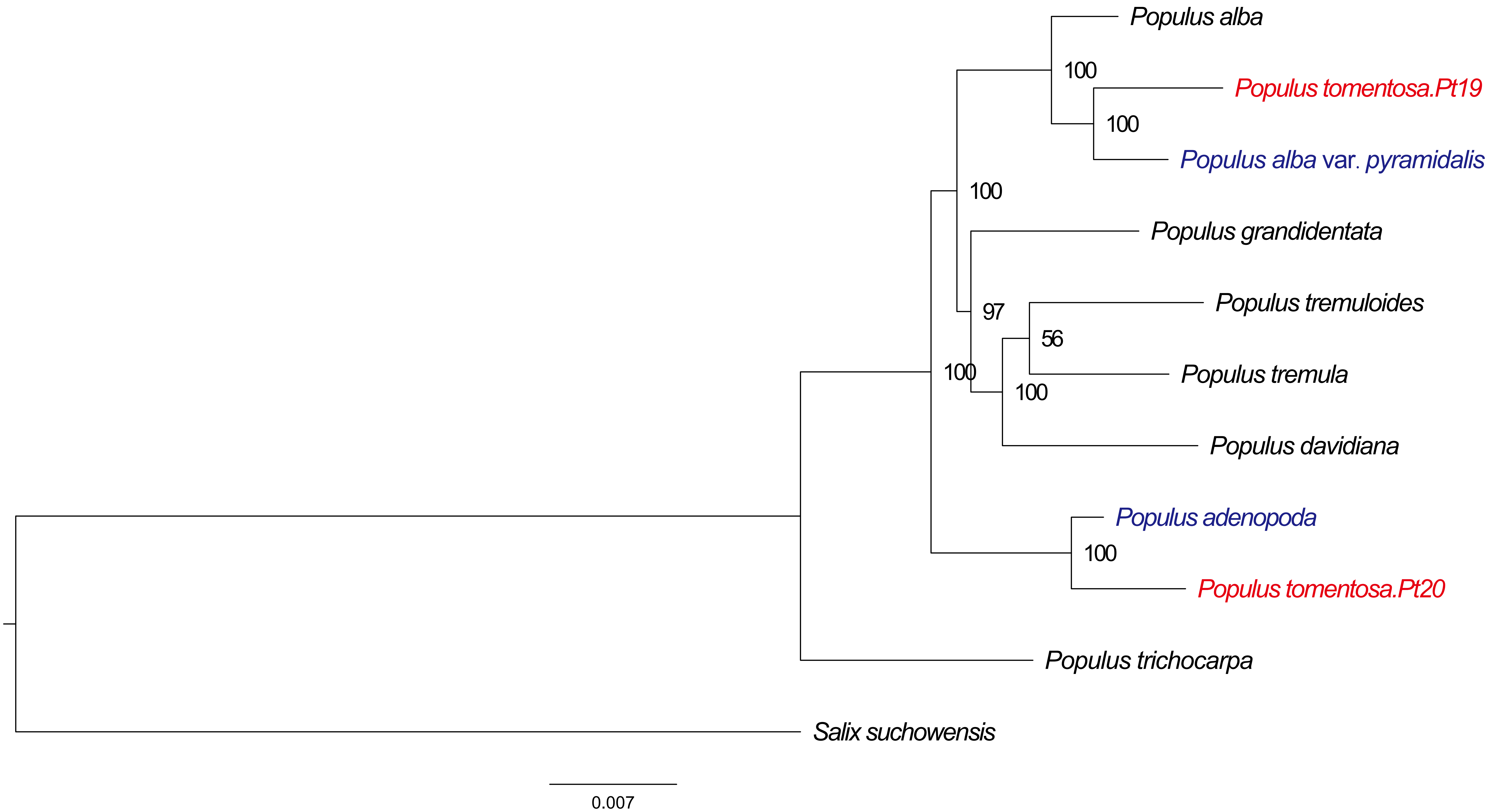
Populus alba
100
Populus tomentosa.Pt19
100
Populus alba var. pyramidalis
100
Populus grandidentata
97
Populus tremuloides
56
Populus tremula
100
100
Populus davidiana
Populus adenopoda
100
Populus tomentosa.Pt20
Populus trichocarpa
Salix suchowensis
0.007

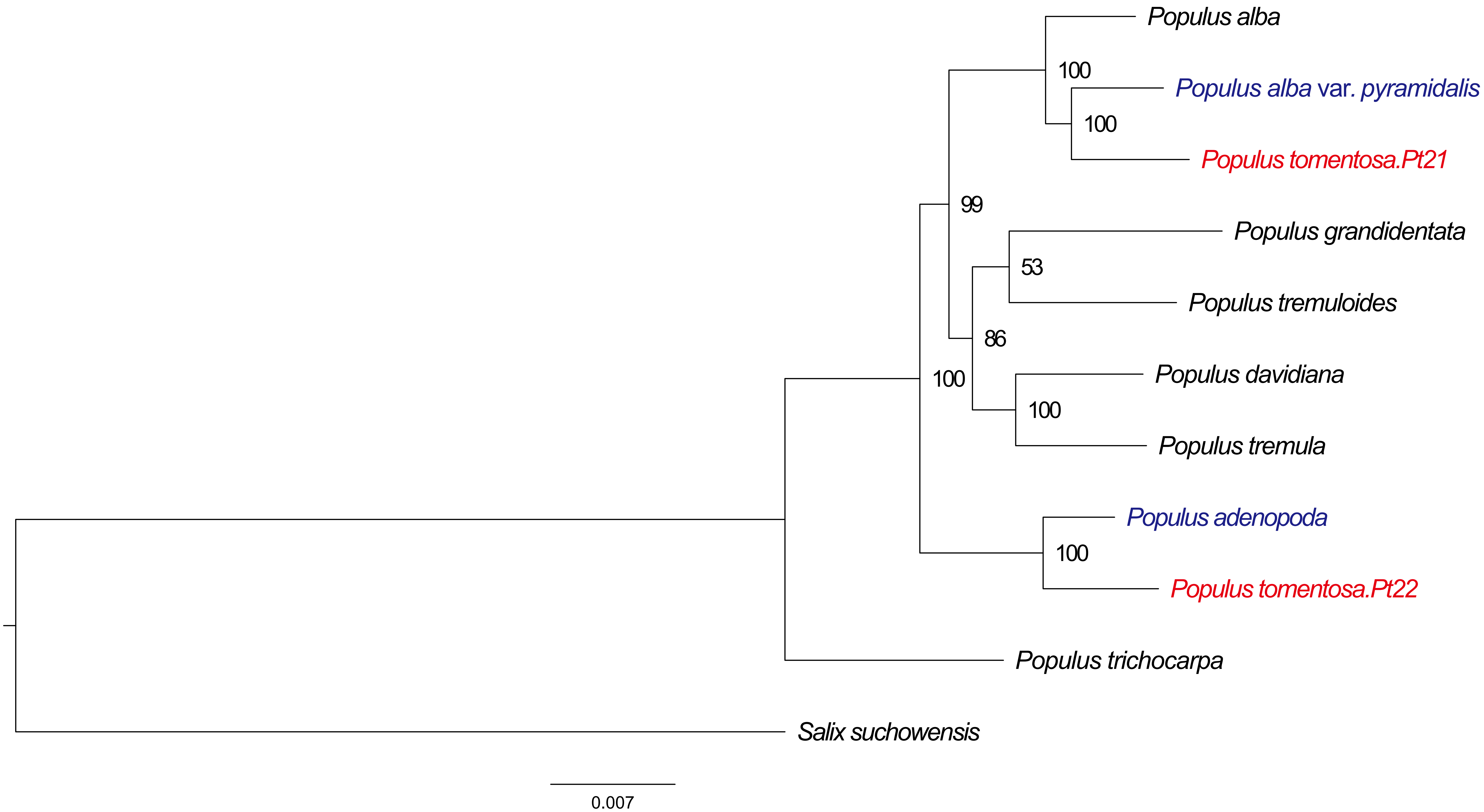

Populus alba
100
Populus alba var. pyramidalis
100
Populus tomentosa.Pt21
99
Populus grandidentata
53
Populus tremuloides
86
Populus davidiana
100
100
Populus tremula
Populus adenopoda
100
Populus tomentosa.Pt22
Populus trichocarpa
Salix suchowensis
0.007

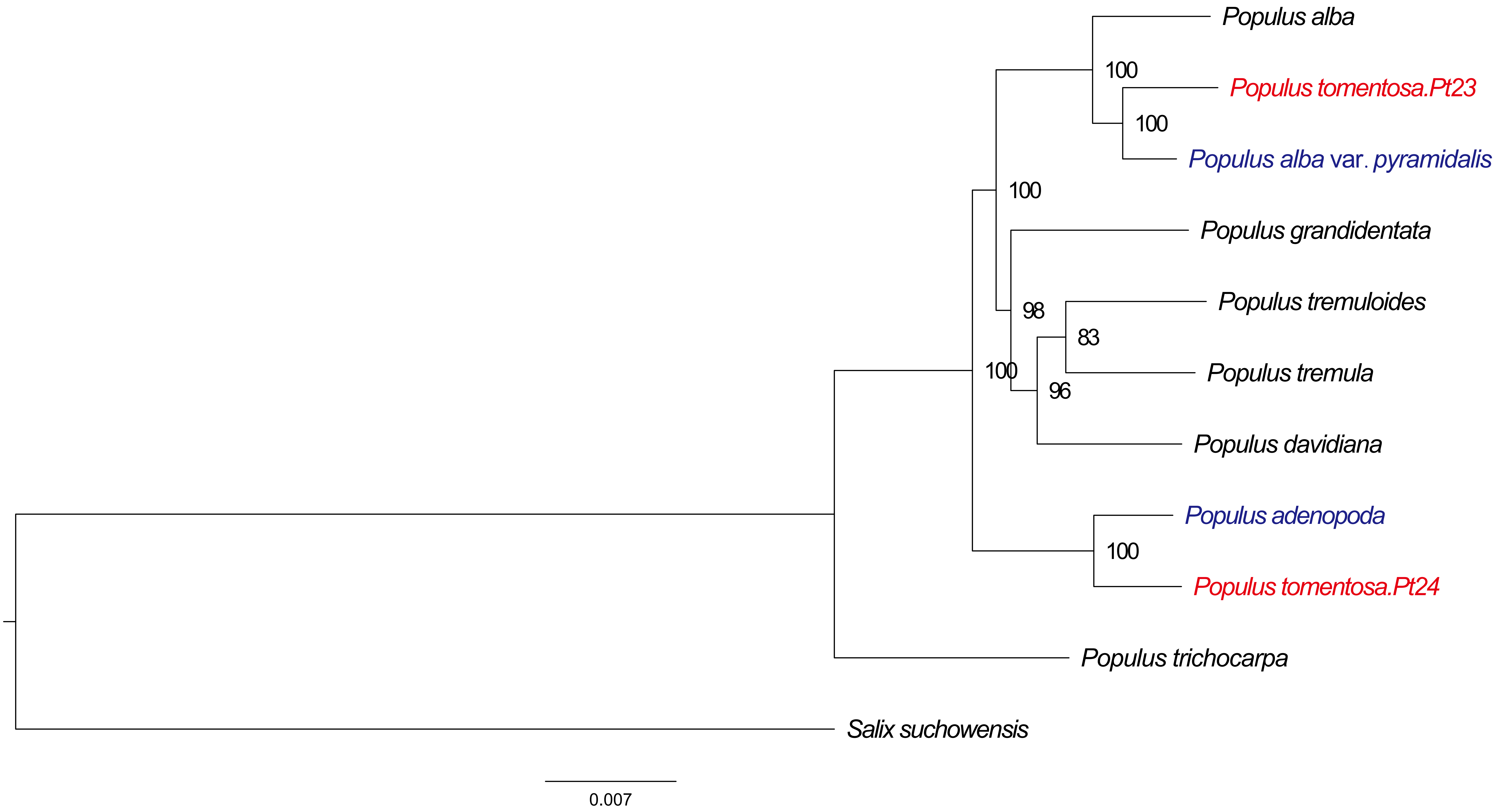
Populus alba
100
Populus tomentosa.Pt23
100
Populus alba var. pyramidalis
100
Populus grandidentata
98
Populus tremuloides
83
Populus tremula
100
96
Populus davidiana
Populus adenopoda
100
Populus tomentosa.Pt24
Populus trichocarpa
Salix suchowensis
0.007

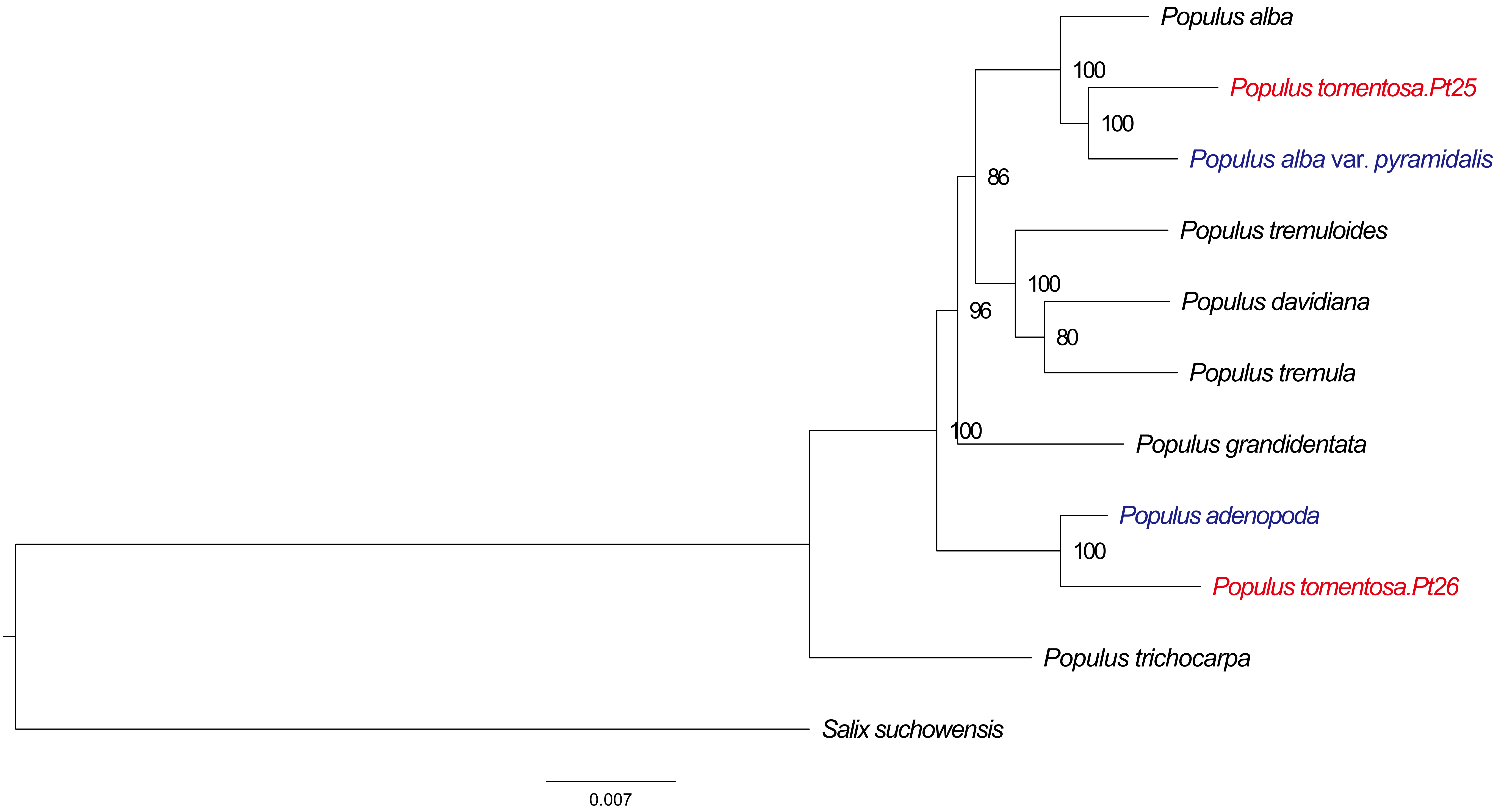

Populus alba
100
Populus tomentosa.Pt25
100
Populus alba var. pyramidalis
86
Populus tremuloides
100
Populus davidiana
96
80
Populus tremula
100
Populus grandidentata
Populus adenopoda
100
Populus tomentosa.Pt26
Populus trichocarpa
Salix suchowensis
0.007

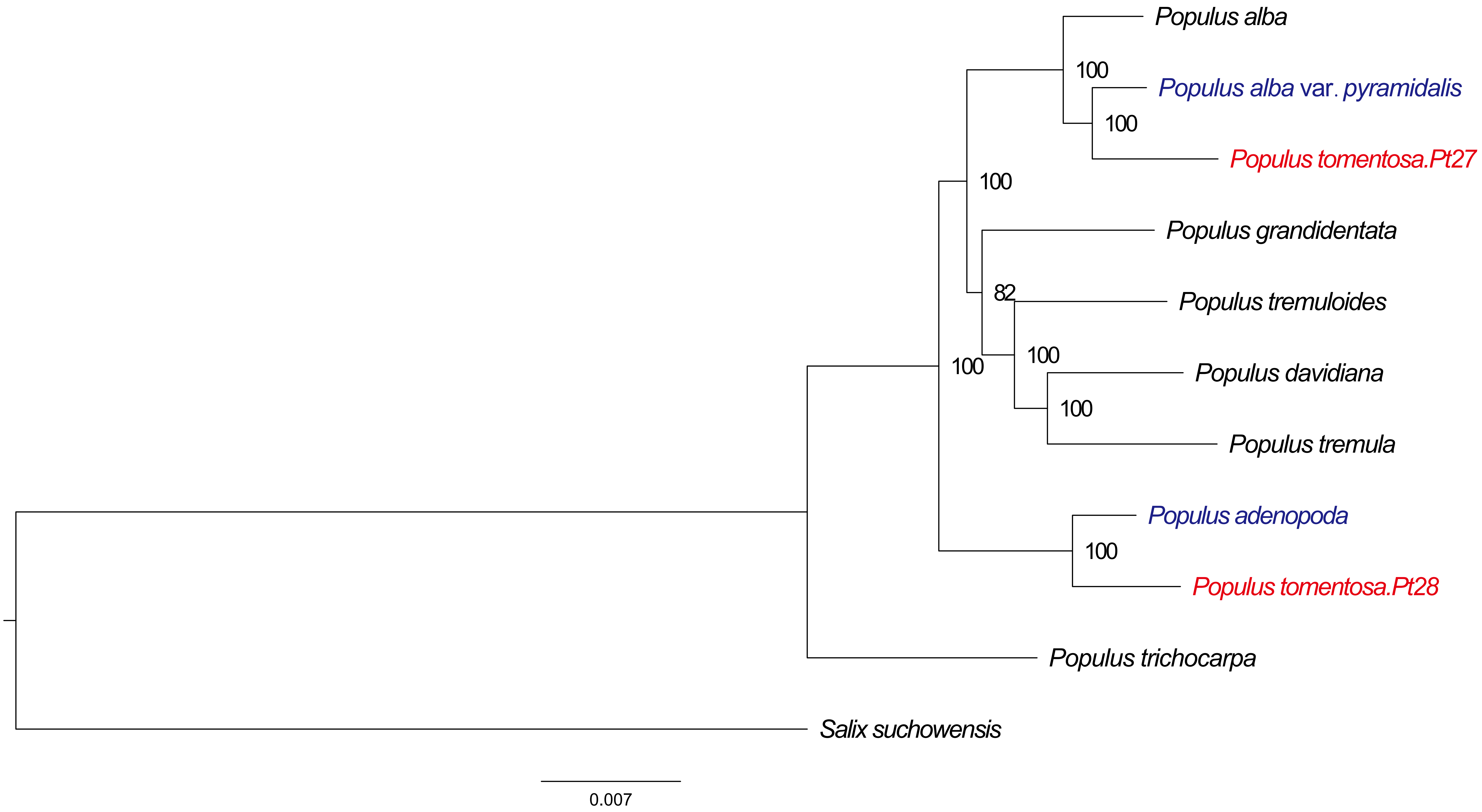

Populus alba
100
Populus alba var. pyramidalis
100
Populus tomentosa.Pt27
100
Populus grandidentata
82
Populus tremuloides
100
Populus davidiana
100
Populus tremula
100
Populus adenopoda
100
Populus tomentosa.Pt28
Populus trichocarpa
Salix suchowensis
0.007

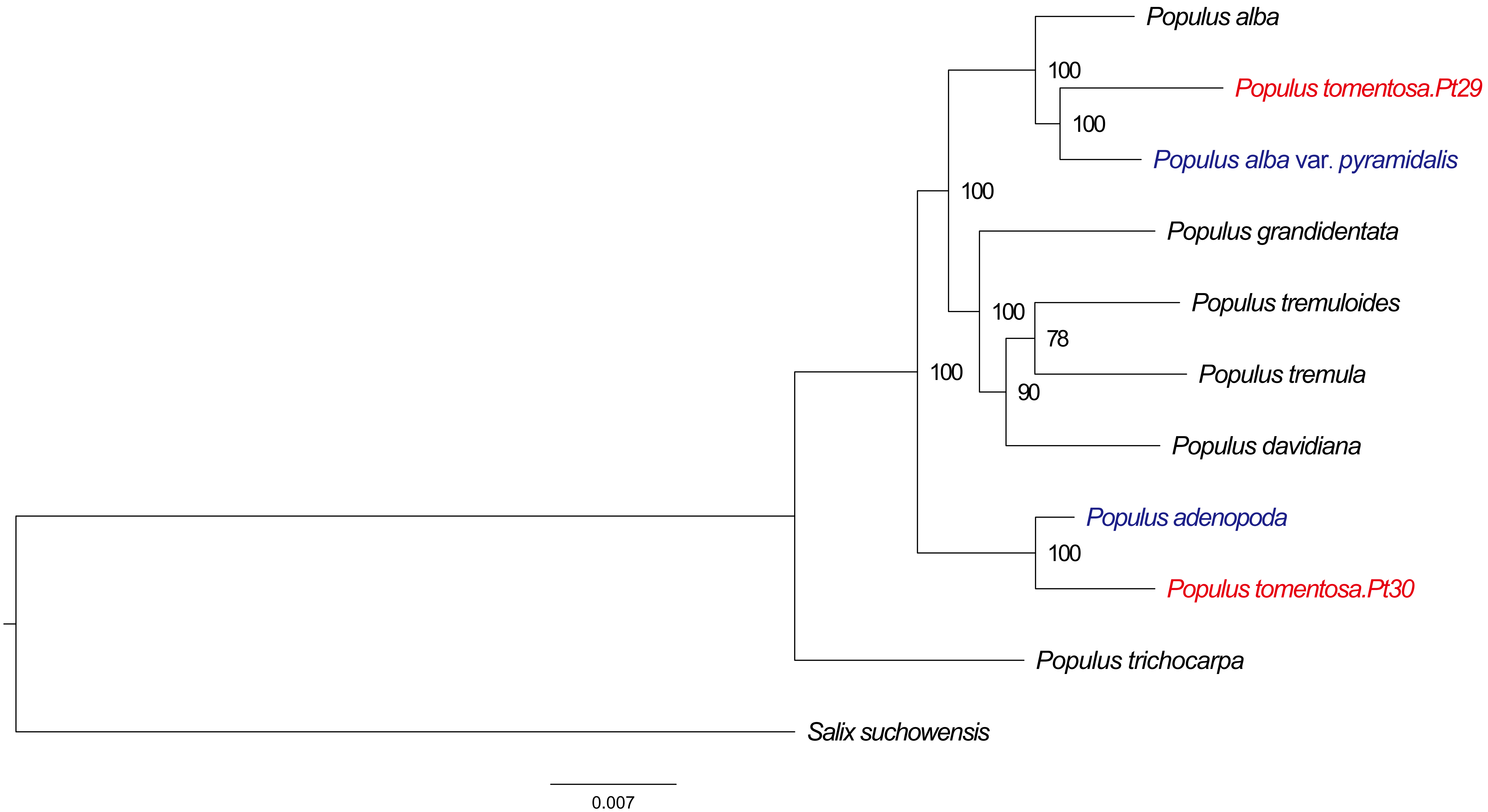
Populus alba
100
Populus tomentosa.Pt29
100
Populus alba var. pyramidalis
100
Populus grandidentata
100
Populus tremuloides
78
Populus tremula
90
100
Populus davidiana
Populus adenopoda
100
Populus tomentosa.Pt30
Populus trichocarpa
Salix suchowensis
0.007

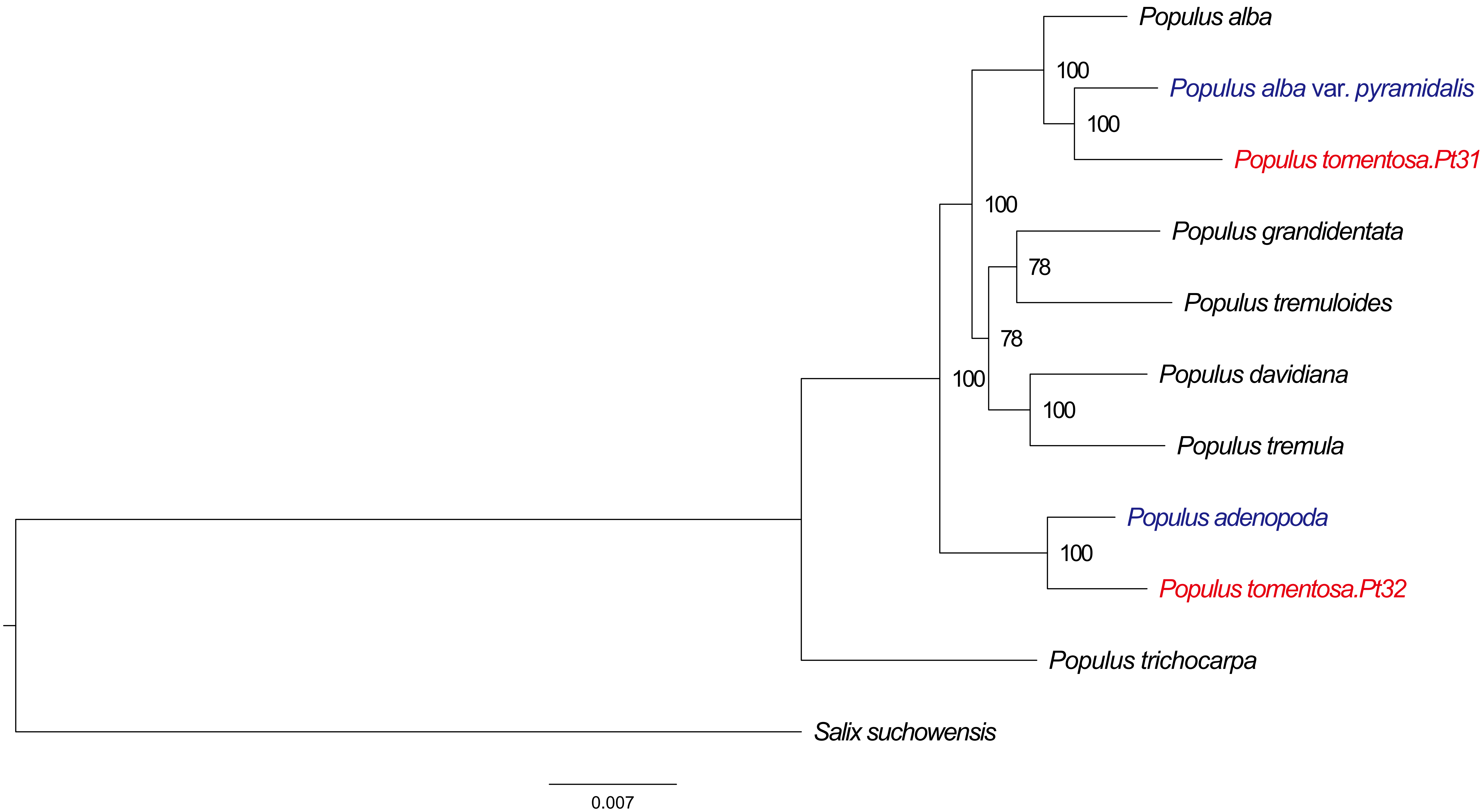
Populus alba
100
Populus alba var. pyramidalis
100
Populus tomentosa.Pt31
100
Populus grandidentata
78
Populus tremuloides
78
100
Populus davidiana
100
Populus tremula
Populus adenopoda
100
Populus tomentosa.Pt32
Populus trichocarpa
Salix suchowensis
0.007

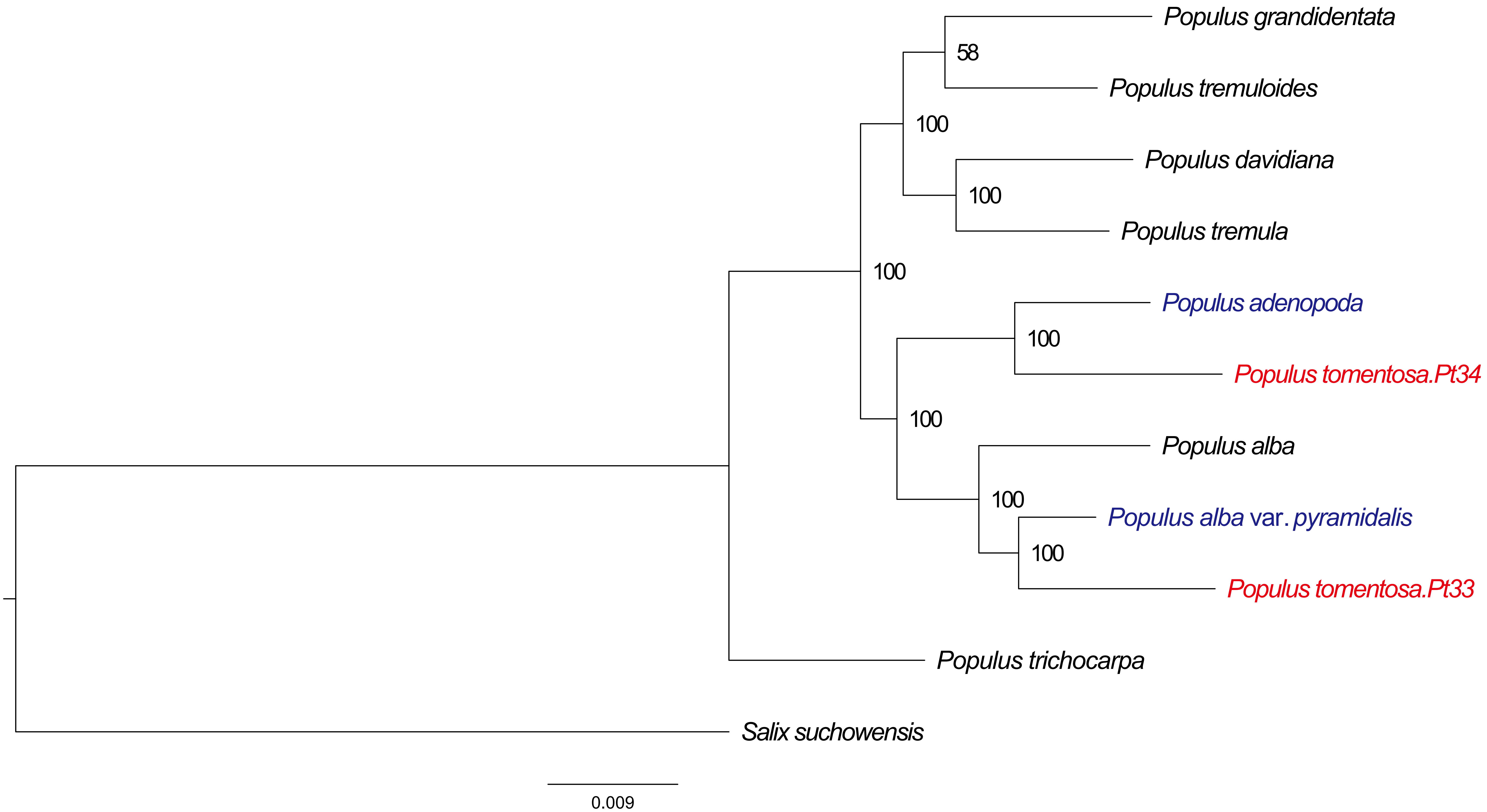
Populus grandidentata
58
Populus tremuloides
100
Populus davidiana
100
Populus tremula
100
Populus adenopoda
100
Populus tomentosa.Pt34
100
Populus alba
100
Populus alba var. pyramidalis
100
Populus tomentosa.Pt33
Populus trichocarpa
Salix suchowensis
0.009

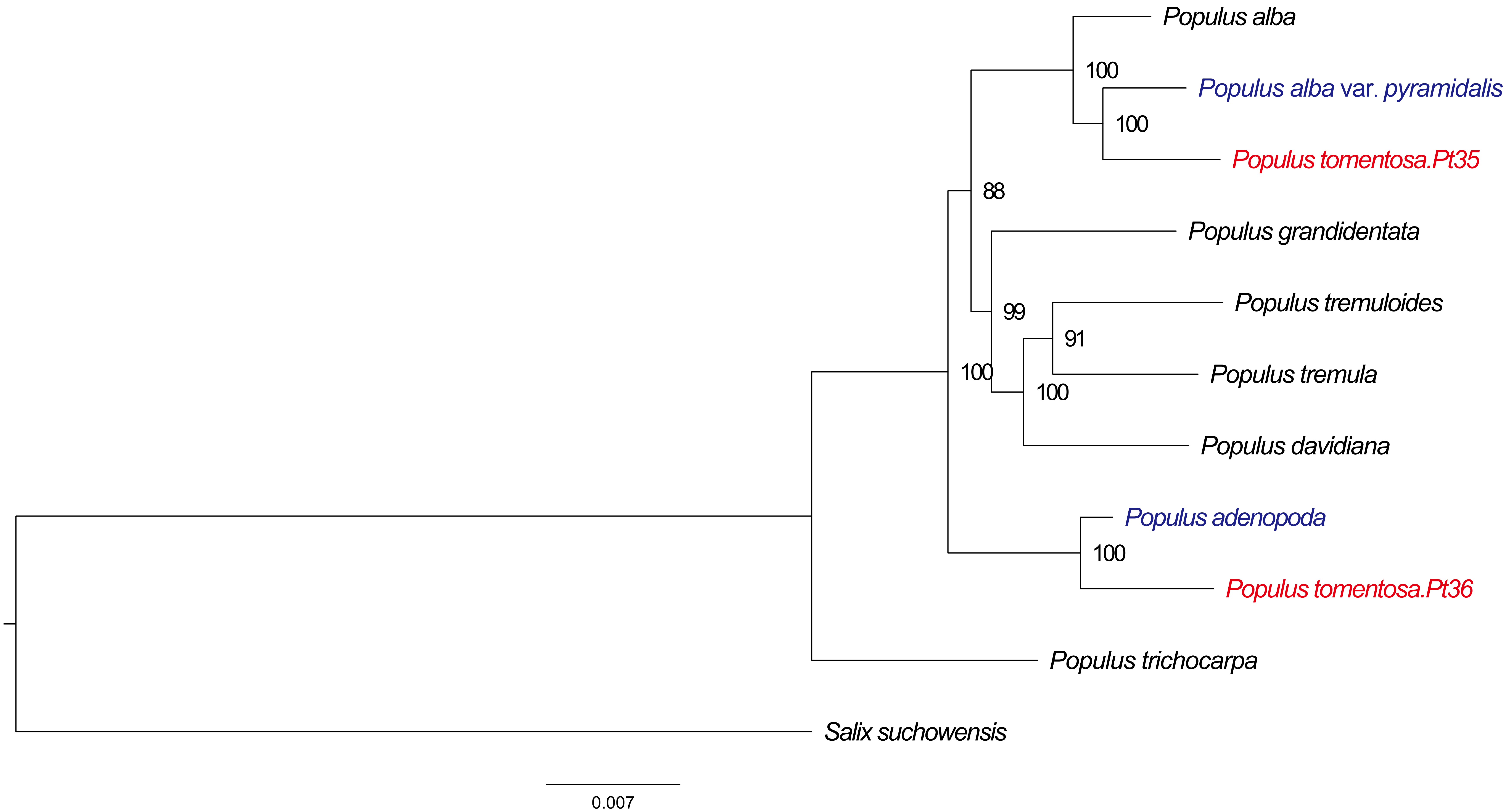
Populus alba
100
Populus alba var. pyramidalis
100
Populus tomentosa.Pt35
88
Populus grandidentata
Populus tremuloides
99
91
Populus tremula
100
100
Populus davidiana
Populus adenopoda
100
Populus tomentosa.Pt36
Populus trichocarpa
Salix suchowensis
0.007

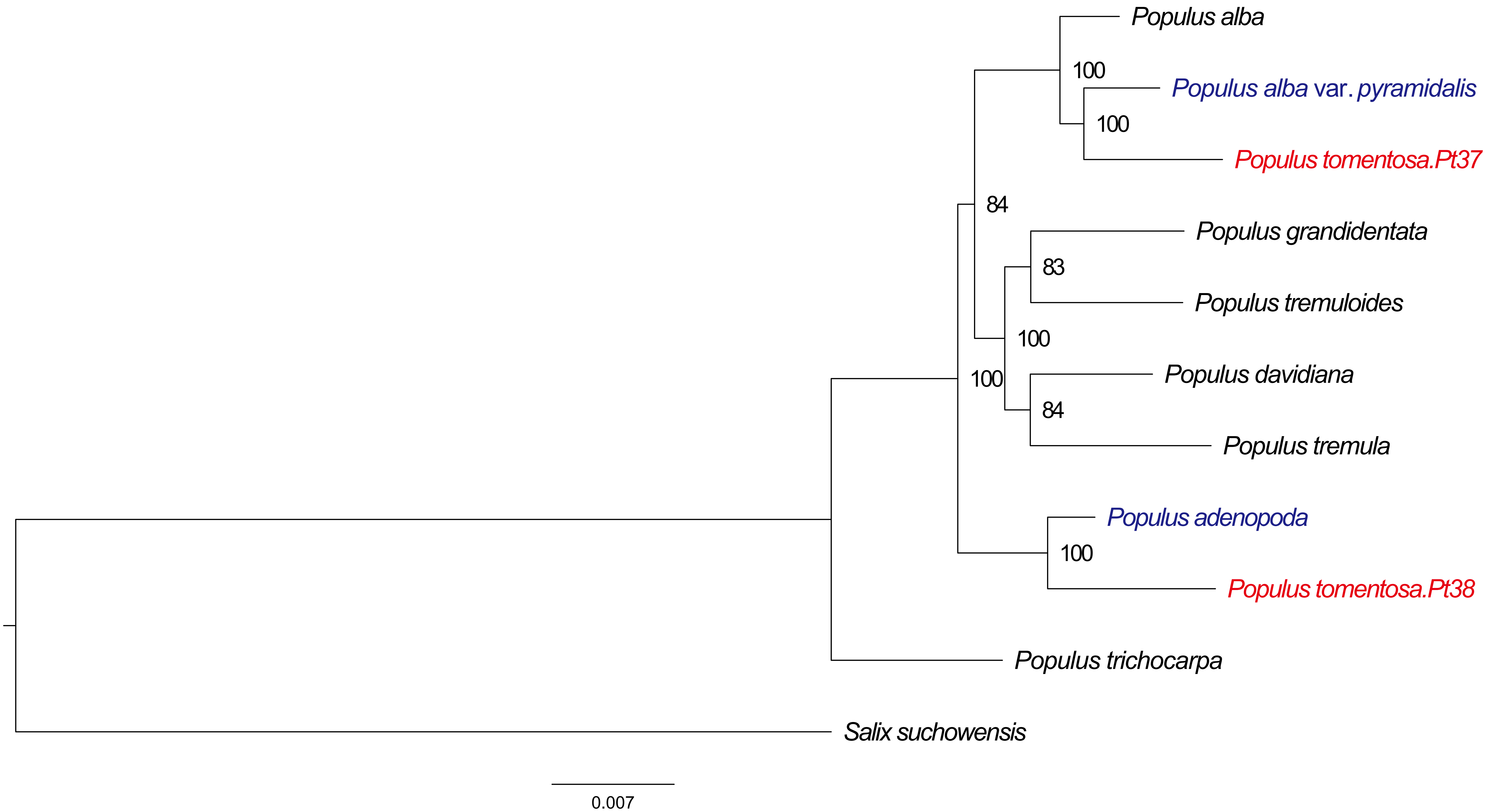
Populus alba
100
Populus alba var. pyramidalis
100
Populus tomentosa.Pt37
84
Populus grandidentata
83
Populus tremuloides
100
100
Populus davidiana
84
Populus tremula
Populus adenopoda
100
Populus tomentosa.Pt38
Populus trichocarpa
Salix suchowensis
0.007
