## Supplementary material for "Hybrid origin of *Populus tomentosa* Carr. identified through genome sequencing and phylogenomic analysis": Genome assembly version and statistics of Populus tomentosa (GM15): Supplementary Table 2 Genome assembly version and statistics.docx

| **Verion** | **Method** | **Genome size** | **Number*** | **N50** | **L50** | **Max.** | **Integrity** (BUSCO^[2]^) |
| --- | --- | --- | --- | --- | --- | --- | --- |
| v0.1 | CANU^[3]^ | 740 Mb | 1300 | 1.94 Mb | 108 | 8.8 Mb | 96.5% |
| v1.0 | v0.1+arrow | 741 Mb | 1300 | 1.94 Mb | 108 | 8.8 Mb | 96.5% |
| v1f | v1.0+arrow×3  +pilon^[7]^×5 | 740 Mb | 4025/2388 | 864 Kb / 34 Mb | 239/8 | 5.5 Mb / 92.4 Mb | 96.5% |

* contig/scaffold

[2]	Simão F A, Waterhouse R M, Ioannidis P et. al. BUSCO: assessing genome assembly and annotation completeness with single-copy orthologs. [J]. Bioinformatics, 2015, 31 (19): 3210

[3]	Koren S, Walenz B P, Berlin K et. al. Canu: scalable and accurate long-read assembly via adaptive k-mer weighting and repeat separation. [J]. Genome Res., 2017, 27 (5): 722

[7]	Walker B J, Abeel T, Shea T et. al. Pilon: An Integrated Tool for Comprehensive Microbial Variant Detection and Genome Assembly Improvement [J]. Plos One, 2014, 9 (11): e112963
