## Supplementary material for "Hybrid origin of *Populus tomentosa* Carr. identified through genome sequencing and phylogenomic analysis": Whole genome of Populus tomentosa (GM15): Supplementary Table 4 Sequencing technology and assembly statistics comparisons of seven poplars genomes.docx

| Genome | Sequencing platform | Assembled genome size | Contigs | | | | Scaffolds | | | | Reference |
| --- | --- | --- | --- | --- | --- | --- | --- | --- | --- | --- | --- |
|  |  |  | Total number | N50 size (kb) | N50 number | Largest (kb) | Total number | N50 size (kb) | N50 number | Largest (kb) |  |
| *P. tomentosa* | PacBio + Hi-C + HiSeq X-10 | 740.2 Mb | 4,025 | 994.5 | 239 | 5,467.9 | 2,407 | 18,914.8 | 38 | 46,677.8 |  |
| *P. trichocarpa* | Sanger+Illumina | 422.9 Mb | NA | 522.8 | 206 | NA | 8,313 | 19,500.0 | 8 | NA | Tuskan et al., 2006 |
| *P. euphratica* | Illumina HiSeq 2000 | 497.0 Mb | 32,882 | 40.4 | 3,078 | 728.0 | 9,673 | 482 | 204 | 8,760 | Ma et al., 2014 |
| *P. pruinosa* | Illumina HiSeq X-10 | 479.3 Mb | 170,219 | 14.0 | NA | 197.6 | 78,960 | 698.5 | NA | 10,688.7 | Yang et al., 2017 |
| *P.alba* var*. pyramidalis* | Illumina + PacBio | 464.0 Mb | 55,988 | 9.8 | NA | 230.3 | 17,797 | 459.2 | NA | 3,427.2 | Ma et al., 2019 |
| *P. alba* | PacBio + HiSeq 2500 | 416.0 Mb | NA | 1,180.9 | 101 | NA | NA | 144.8 | 473 | NA | Liu et al., 2019 |
| 84K poplar | PacBio + Hi-C + Illumina | 747.5 Mb | NA | 1,990.0 | NA | NA | NA | 19,600.0 | NA | NA | Qiu et al., 2019 |

NA: not available
