## Supplementary material for "Hybrid origin of *Populus tomentosa* Carr. identified through genome sequencing and phylogenomic analysis": Gene annotation of Populus tomentosa (GM15): Supplementary Table 8. Gene annotation of Populus tomentosa (GM15).docx

| **Genes** | **Swiss_Prot** | **TrEMBL** | **NR** | **Pfam** | **KOG** | **GO** | **KO** |
| --- | --- | --- | --- | --- | --- | --- | --- |
| 59,124 | 37,675 | 58,175 | 58,276 | 49,743 | 56,191 | 47,205 | 22,575 |
| 100.00% | 63.70% | 98.40% | 98.60% | 84.10% | 95.00% | 79.80% | 38.20% |
