## Supplementary material for "Hybrid origin of *Populus tomentosa* Carr. identified through genome sequencing and phylogenomic analysis": Cafe summary-tree of Populus tomentosa(GM15) and other species: Supplementary Table 10 Recombination test results.docx

Supplementary Table 10. Recombination analysis between two subgenomes of *P.tomentosa*

|  |  | Percent % | Percent % |
| --- | --- | --- | --- |
| Total test loci | 5345 |  |  |
| Non-recombination | 4309 | 80.62% | 99.13% |
| Recombination | 38 | 0.71% | 0.87% |
| Not meet the above two conditions | 998 | 18.67% |  |
